# Proximity Labeling to Map Host-Pathogen Interactions at the Membrane of a Bacteria Containing Vacuole in *Chlamydia trachomatis* Infected Human Cells

**DOI:** 10.1101/616896

**Authors:** Macy G. Olson, Ray E. Widner, Lisa M. Jorgenson, Alyssa Lawrence, Dragana Lagundzin, Nicholas T. Woods, Scot P. Ouellette, Elizabeth A. Rucks

## Abstract

As an obligate intracellular pathogenic bacterium, *C. trachomatis* develops within a membrane-bound vacuole, termed the inclusion. The inclusion membrane is modified by chlamydial inclusion membrane proteins (Incs), which act as the mediators of host-pathogen interactions. An *in vivo* understanding of Inc-Inc and Inc-eukaryotic protein interactions and how these contribute to overall host-chlamydial interactions at this unique membrane is lacking. Previous bacterial two-hybrid studies established that certain Incs have the propensity to bind other Incs while others have limited Inc-Inc interactions. We hypothesize some Incs organize the inclusion membrane whereas other Incs bind eukaryotic proteins to promote chlamydial-host interactions. To test this hypothesis, we used the ascorbate peroxidase proximity labeling system (APEX2), which labels proximal proteins with biotin *in vivo*, and chose to analyze Inc proteins with varying Inc-binding propensities. We inducibly expressed these Incs fused to APEX2 in *Chlamydia trachomatis* L2, verified their localization and labeling activities by transmission electron microscopy, and used affinity purification-mass spectrometry to identify biotinylated proteins. To analyze our mass spectrometry results for statistical significance, we used Significance Analysis of INTeractome (SAINT), which demonstrated that our Inc-APEX2 constructs labeled Inc proteins as well as known and previously unreported eukaryotic proteins that localize to the inclusion. Our results broadly support two types of Inc interactions: Inc-Inc versus Inc-host. One eukaryotic protein, LRRFIP1 (LRRF1) was found in all of our Inc-APEX2 datasets, which is consistent with previously published AP-MS datasets. For the first time, we demonstrate by confocal and super-resolution microscopy that endogenous LRRF1 localizes to the chlamydial inclusion. We also used bacterial two-hybrid studies and pulldown assays to determine if LRRF1 was identified as a true interacting protein or was proximal to our Inc-APEX2 constructs. Combined, our data highlight the utility of APEX2 to capture the complex *in vivo* protein-protein interactions at the chlamydial inclusion.

**Author summary:** Many intracellular bacteria, including the obligate intracellular pathogen *Chlamydia trachomatis*, grow within a membrane-bound “bacteria containing vacuole” (BCV) that, in most cases, prevents association with the lysosome. Secreted cytosolic effectors modulate host activity, but an understanding of the host-pathogen interactions that occur at the BCV membrane is limited by the difficulty in purifying membrane fractions from infected host cells. Here, we used the ascorbate peroxidase proximity labeling system (APEX2), which labels proximal proteins with biotin *in vivo*, to study the interactions that occur at the chlamydial vacuolar, or inclusion, membrane. The inclusion membrane is modified by chlamydial type III secreted inclusion membrane proteins (Incs), which act as the mediators of host-pathogen interactions. Our results broadly support two types of Inc interactions: Inc-Inc versus Inc-host. Our data highlight the utility of APEX2 to capture the complex protein-protein interactions at a membrane site *in vivo* in the context of infection.

## Introduction

*Chlamydia trachomatis* is the leading cause of bacterial sexually transmitted infections (1). In 2017, 1.7 million cases were reported in the United States, with the highest incidence of infection in people ages 15 to 29 (2). Approximately 75% of infections are asymptomatic, and prolonged infection in women can lead to pelvic inflammatory disease and ectopic pregnancy (1). Infections in men can cause urethritis, epididymitis, and prostatitis (3, 4). Asymptomatic infections likely occur due to the obligate intracellular nature of this pathogen and manipulation of host cell responses by chlamydial secreted effectors (1).

*Chlamydiae* are developmentally regulated pathogens that reside within a membrane-bound vacuole, called an inclusion. *C. trachomatis* has two developmental forms: the infectious elementary body (EB) and the non-infectious, reticulate body (RB). The EB infects a host cell, differentiates into an RB, and develops within a membrane-bound vacuole, termed an inclusion. The inclusion is initially derived from the eukaryotic plasma membrane that engulfs the invading EB and forms a barrier between the host and pathogen (1, 5). Within the first few hours of infection, the chlamydial inclusion disassociates from the endosomal/lysosomal pathway. This process is likely mediated by the active modification by *Chlamydia* of the inclusion membrane via the insertion of type III secreted chlamydial inclusion membrane proteins (Incs) (6) and the recruitment of lipids and other host proteins to the chlamydial inclusion (7-15). Incs contain two or more hydrophobic transmembrane domains with both termini located on the host cytosolic face of the inclusion (5, 16-18). An estimated 50-70 *inc* genes (19) account for approximately seven percent of the highly reduced chlamydial genome indicating that these genes are important for optimal chlamydial development (20). In addition, Incs are temporally expressed throughout the developmental cycle (17, 21-23), which suggests that there are likely dedicated roles at specific points during the developmental cycle for individual Incs in the inclusion membrane.

To maximize the production of infectious EB progeny, *C. trachomatis* must recruit the necessary nutrients it needs to develop yet protect against the host immune response. Given that the inclusion membrane is the host-pathogen interface and that *Chlamydiae* extensively modify this membrane with secreted Incs, Inc proteins are likely central to achieve these functions. We hypothesize that Incs serve two functions: (i) to organize the inclusion membrane by forming nodes of interaction and spatially coordinating Inc-Inc interactions and (ii) to recruit eukaryotic proteins to facilitate necessary host-chlamydial interactions. Both functions are important to complete the developmental cycle and likely are not mutually exclusive. In support of this hypothesis, a previous bacterial two-hybrid (BACTH) study indicated that specific Inc proteins (e.g., IncF) bind multiple Incs (23), while other Incs (e.g., IncA) have been shown to interact with eukaryotic proteins (9, 12, 14, 24-26). Previous work has shown that knocking out certain Incs results in a weakened inclusion membrane and premature lysis (27). Although Incs represent the vast majority of identified chlamydial type III secreted proteins, little is known about their function in the inclusion membrane. This is largely due to the inherent difficulties of purifying Incs that contain large hydrophobic regions (5), where the conditions required for solubilization do not preserve protein-protein interactions. By identifying the totality of protein binding partners for Incs during chlamydial infection, the function of specific Inc-protein interactions at the inclusion can be understood.

Until recently, *C. trachomatis* was genetically intractable, which had been a major hindrance in advancing *C. trachomatis*-host interaction research. In the past, *in vitro* methods were used to identify Inc-protein binding partners. Such methods included transient transfection of epitope-tagged chlamydial Incs in uninfected host cells or running whole cell uninfected lysates over a column bound by the soluble domain of a recombinant Inc protein (24). One drawback of ectopic expression is that the Inc proteins are expressed in the host cell out of their normal spatial context (e.g., they tend to aggregate in micelle-type structures) (28), increasing the possibility of identifying false interactions. Moreover, the ectopic expression of Inc proteins in eukaryotic cells, in contrast to their type III secretion from *Chlamydiae*, is unlikely to result in correct protein folding and subcellular location (i.e. in the inclusion membrane). A recombinant Inc bound to a column and exposed to total eukaryotic cell lysate can promote false positive interactions between eukaryotic proteins that are in subcellular compartments that do not typically interact with the chlamydial inclusion. An alternative strategy to purify chlamydial inclusions from large numbers of infected host cells requires lysis conditions and density gradient purification steps that can disrupt transient protein-protein interactions at the inclusion membrane and often results in lysis of the fragile inclusion membrane, yielding a total recovery rate of only about eight percent (29). Furthermore, this purification method is labor-intense and requires equipment that might not be available to other laboratories (29). In addition, and equally important, such methods do not capture potential Inc-Inc interactions. Not surprisingly, the Inc protein binding partners identified to date using these methods are eukaryotic proteins. For example, IncG has been shown to bind the eukaryotic protein 14-3-3β to modify host signaling (9), IncD binds ceramide transport protein, CERT, to acquire lipids (12), and IncE binds sorting nexin (SNX) 5 and SNX6 to interfere with cargo trafficking (24, 25).

To capture dyanamic *in vivo* protein-protein interactions, including Inc-Inc interactions at the chlamydial inclusion, we were the first to use and characterize the feasibility, including important caveats, of using the ascorbate peroxidase proximity labeling system (APEX2) to identify Inc binding partners in the context of *C. trachomatis* infection (30). APEX2, a mutated soybean peroxidase (31, 32), can be fused to a protein of interest and activated during a short (one minute) reaction to covalently modify proximal proteins with a biotin molecule (32). This system can be used to capture *in vivo* “snapshots” of the dynamic protein-protein interactions that occur at the chlamydial inclusion during development. Incs fused to APEX2 are secreted by *C. trachomatis* and inserted in the inclusion membrane (30). Proteins proximal to the expressed Inc-APEX2 fusion protein are covalently modified with biotin after the addition of biotin-phenol and hydrogen peroxide to catalyze the APEX2 biotinylation reaction (30). An additional advantage of using APEX2 to identify Inc protein binding partners is the ability to use high concentrations of detergent to solubilize hydrophobic membrane proteins, like Incs, because there is no need to maintain binding partners after the covalent addition of biotin to neighboring proteins (30). Subsequently, the cells are lysed, and the biotinylated proteins are affinity purified using streptavidin beads and identified using mass spectrometry (AP-MS). A recent publication has highlighted the APEX2 proximity labeling system (33), but their conclusions relied on the use of (i) statistical analyses that fail to adequately account for background contaminants (34), (ii) an Inc whose expression did not resemble that of its endogenous form and did not identify its known interacting Incs (28), and (iii) early time points that did not take into account mitochondrial background contaminants (35, 36).

We applied APEX2 to test our hypothesis using two Incs, IncF and IncA. IncF may be primarily involved in organizing the inclusion because it has been shown to interact extensively with other Incs via BACTH studies (23) and is expressed early after infection (22). IncA may primarily interact with eukaryotic proteins as it contains a eukaryotic SNARE-like domain (37) and has been shown to bind fewer Incs by the same BACTH studies (23). We also created a truncated IncA (IncA_TM_) to interrogate if removal of the C-terminal domain of IncA would alter the specificity of proteins labeled by this construct (30). Using the APEX2 proximity labeling system, we tested these interactions *in vivo* by transforming *C. trachomatis* serovar L2 with Inc-APEX2 fusion constructs that localize to the inclusion membrane when expressed (30). These experiments have helped define the function of Incs in the inclusion membrane and whether Incs collaborate to support chlamydial development within the inclusion.

Our AP-MS studies were analyzed for statistical significance using a Bayesian-based statistical analysis tool, Significance Analysis of INTeractome (SAINT) (38). In each Inc-APEX2 dataset, we identified chlamydial Inc proteins that were statistically significant, and we also identified eukaryotic proteins that had been previously shown to localize with the chlamydial inclusion. Importantly, we identified previously undescribed eukaryotic proteins at the inclusion membrane. Leucine-Rich Repeat Flightless-Interacting Protein 1 (LRRF1) was identified in all of our Inc-APEX2 datasets and has been identified in other AP-MS studies (24, 29, 33). We also identified a LRRF1 binding partner, Protein Flightless 1 homolog (FLI1), in our IncA-APEX2 dataset, indicating that we were also identifying partial signaling pathways.

The presence of LRRF1 in our datasets gave us the opportunity to ask why this is a prominently identified protein in ours and others’ AP-MS studies. We were skeptical that one protein was a true interactor with every single one of our Inc-APEX2 constructs. Therefore, we designed a series of experiments to help us understand how LRRF1 was identified, either through direct interaction with one of our Inc-APEX2 constructs or interaction with an adjacent Inc, but within the labeling radius of our Inc-APEX2 constructs. For the first time, we demonstrate that endogenous LRRF1 and FLI1 localize with the chlamydial inclusion. LRRF1 localization with the inclusion was conserved between closely related *C. trachomatis* serovars and strains. By bacterial two-hybrid assay, LRRF1 was found to interact with the Inc, CT226, which is consistent with a previous study which identified LRRF1 and FLI1 by transfecting Strep-tagged CT226 into uninfected eukaryotic cells (24). We also performed a co-immunoprecipitation using CT226_FLAG_ expressed from *C. trachomatis* and identified LRRF1 in the eluate. Overall, our proximity labeling system has identified both known and previously unreported proteins at the inclusion membrane and highlights the utility of an *in vivo* proximity labeling system to identify protein-protein interactions and how they are recruited to the chlamydial inclusion membrane.

## Results

### Biotinylation of proximal proteins at the inclusion membrane using *C. trachomatis* L2 Inc-APEX2 transformants

To examine our hypothesis that some Incs preferentially interact with other Inc proteins whereas other Incs primarily interact with eukaryotic proteins, we used the ascorbate peroxidase proximity labeling system (APEX2) to determine chlamydial Inc binding partners *in vivo* (30). To do this, we transformed *C. trachomatis* serovar L2 with a plasmid encoding IncF-APEX2, IncA_Transmembrane Domain_-APEX2 (IncA_TM_-APEX2), IncA-APEX2, or APEX2 only controlled by an anhydrotetracycline (aTc) inducible promoter system. IncF has previously been shown to interact with several Incs (23) and contains a short cytosolic domain, which could limit its ability to interact with eukaryotic proteins. In the same study, IncA interacted with fewer Incs (23) and has a large cytosolic domain with a eukaryotic SNARE-like domain (37, 39-41), suggesting that IncA might preferentially interact with eukaryotic proteins. IncA_TM_-APEX2 is truncated to have a short cytosolic domain like IncF-APEX2 and is to determine if the C-terminus of IncA confers specificity towards determining protein binding partners (30). APEX2 only is a negative control included in each experiment and, when expressed in transformed *C. trachomatis* L2, remains in the bacterial cytosol because it lacks the type III secretion signal (Fig 1) (30). All proximity labeling experiments were performed using a plaque-cloned population of *C. trachomatis* L2 transformants. HeLa 229 cells were infected with *C. trachomatis* L2 IncF-APEX2, IncA_TM_-APEX2, IncA-APEX2, or APEX2 transformants and induced with anhydrotetracycline (aTc). As previously determined, this resulted in the expression and localization of each construct that matches endogenous IncA and IncF (21, 30, 42). An epitope tag (FLAG) is located at the N-terminus of APEX2 and is used to visualize the localization of the various APEX2 constructs expressed from *C. trachomatis* L2.

**Figure 1.**
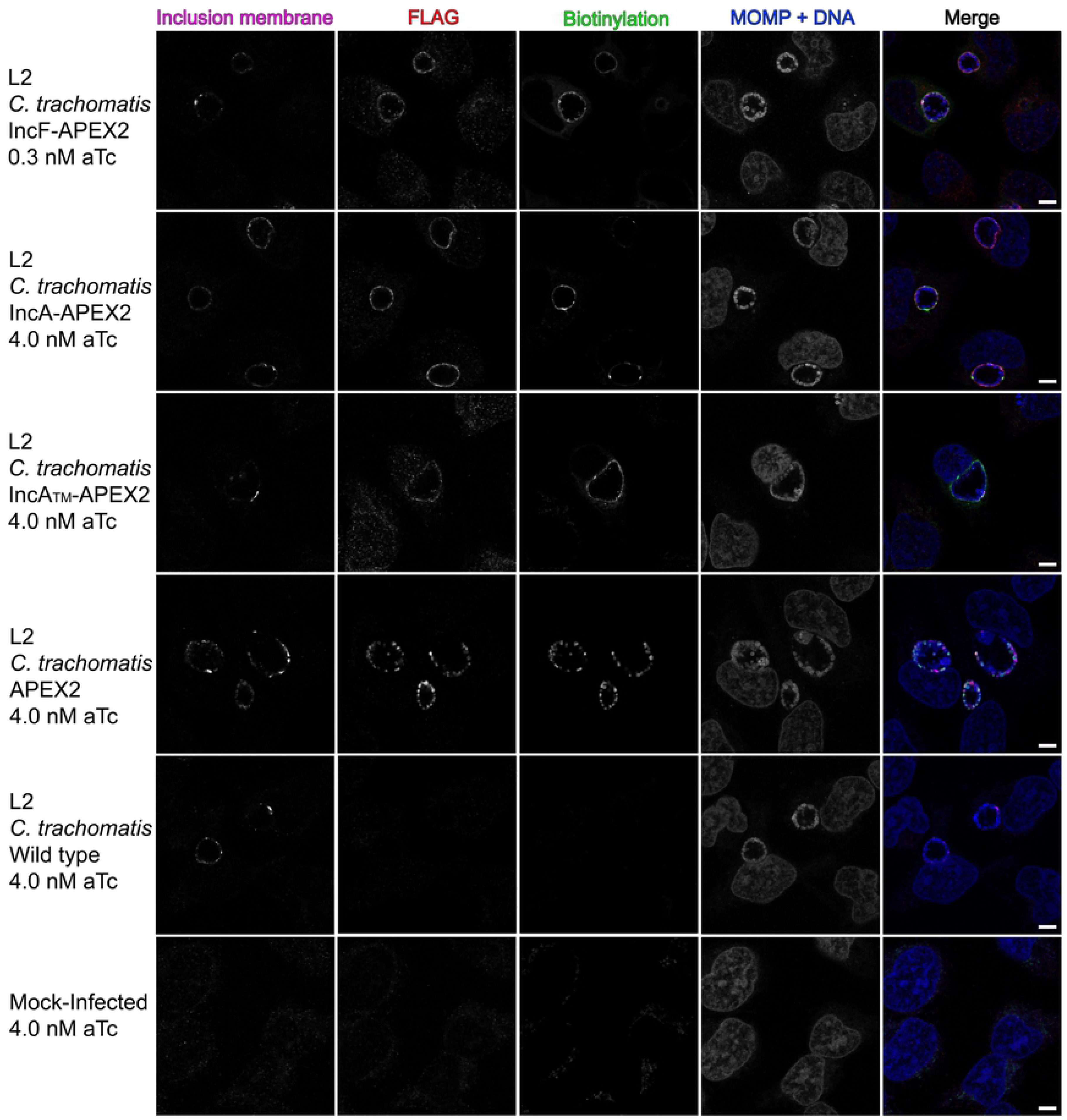
Localization and biotinylation of proteins proximal to the inclusion membrane in HeLa cells infected with *C. trachomatis* L2 transformants expressing Inc-APEX2 constructs. Coverslips were placed in two wells of a 6-well tissue culture plate to ensure appropriate biotinylation. HeLa cells infected with *C. trachomatis* serovar L2 transformed with the indicated APEX2 constructs, *C. trachomatis* L2 wild-type (WT), or mock-infected were induced for construct expression with the indicated concentrations of anhydrotetracycline (aTc) at 7 hpi. Biotin-phenol was added at 23.5 hpi, biotinylation was catalyzed at 24 hpi by the addition of 3 mM H_2_O_2_ for 1 min, after which the reaction was quenched. Coverslips were removed from the 6-well plate and processed for immunofluorescence to visualize biotinylated proteins (streptavidin-488 conjugate), expression of the construct (anti-Flag, red), *Chlamydiae* (MOMP) and DNA (DAPI; blue), and the inclusion membrane (anti-CT223, pink). Coverslips were imaged using a Zeiss confocal LSM 800 with 63x magnification and 2x zoom. Scale bar = 5 µm.

For each biotinylation experiment, coverslips were placed in two wells of a six-well plate to confirm the presence of biotinylated proteins at the inclusion membrane by indirect immunofluorescence microscopy. This was performed for each of the test conditions and controls. HeLa cells were infected with *C. trachomatis* L2 IncF-APEX2, IncA_TM_-APEX2, IncA-APEX2, or APEX2 and construct expression was induced at 7 hpi (0.3 nM aTc for IncF-APEX2 and 4 nM aTc for all other transformants). Biotin-phenol was added to each well at 23.5 hpi, and at 24 hpi hydrogen peroxide (H_2_O_2_) was added to the wells to catalyze the biotinylation reaction during a one-minute incubation. After biotinylation, the enzymatic APEX2 activity was quenched. Coverslips were removed from the wells, fixed, then stained for immunofluorescence to confirm appropriate biotinylation. Separately, the lysate was collected as indicated in the *Methods* and processed after confirming biotinylation by indirect immunofluorescence. Expression of each construct containing APEX2 and biotinylation at the inclusion membrane was observed using each of the *C. trachomatis* Inc-APEX2 transformants (Fig 1). For *C. trachomatis* L2 APEX2, which lacks a type III secretion signal, biotinylation was co-localized with the bacterial cytosol (Fig 1). For *C. trachomatis* L2 wild-type (i.e., untransformed) infected and mock-infected HeLa cells (i.e., no APEX2), faint biotinylation was observed in subcellular structures consistent with mitochondria, but no biotinylation was detected at the inclusion of *C. trachomatis* L2 wild-type (Fig 1). This confirmed that proteins proximal to the inclusion were biotinylated using the Inc-APEX2 constructs.

### Verification of Inc-APEX2 labeling activity on the cytosolic face of the inclusion membrane by electron microscopy

*C. trachomatis* L2 transformed with IncF-APEX2, IncA_TM_-APEX2, and IncA-APEX2 target the constructs to the inclusion membrane with the C-terminus (that contains APEX2) exposed to the host cytosol (43). We used electron microscopy to further support that the *C. trachomatis* L2 Inc-APEX2 transformants were labeling the cytosolic face of the chlamydial inclusion (44). For these studies, HeLa cells were infected with wild-type (i.e., untransformed) or *C. trachomatis* L2 Inc-APEX2 transformants, and monolayers were treated with aTc to induce APEX2 fusion protein expression. Then, cells were fixed with a glutaraldehyde and paraformaldehyde solution, which maintains APEX2 activity (44), and labeled with or without 3,3′-Diaminobenzidine (DAB). DAB and hydrogen peroxide (H_2_O_2_) diffuse into non-permeabilized cells, and, in the proximity of APEX2, DAB polymerizes (31, 44, 45). Upon polymerization, DAB becomes membrane impermeable and remains closely associated with the site of polymerization (44). DAB reacts with the heavy metal staining procedure (osmium tetroxide) to create a contrast that can be observed by electron microscopy (44).

As seen in Fig 2A, no DAB polymerization is observed at the inclusion membrane in HeLa cells infected with wild-type *C. trachomatis* L2. To control for background activity, HeLa cells were infected with *C. trachomatis* L2 IncA-APEX2, induced for expression, but not treated with DAB (Fig 2B). In these samples, no DAB staining was observed at the inclusion membrane (Fig 2B). There was no detectable DAB labeling at the inclusion membrane in HeLa cells infected with *C. trachomatis* L2 APEX2, but we did not observe strong DAB polymerization within individual organisms (Fig 2C). In HeLa cells infected with *C. trachomatis* L2 IncF-APEX2, IncA_TM_-APEX2, or IncA-APEX2 transformants, DAB polymerization was observed at the inclusion membrane (Fig 2C, arrows). However, there appeared to be less DAB labeling with IncA_TM_-APEX2 compared to IncF-APEX2 and IncA-APEX2. Overall, by electron microscopy, we observed Inc-APEX2 directed DAB labeling at the inclusion membrane, and this labeling appeared on the cytosolic face of the inclusion membrane.

**Figure 2.**
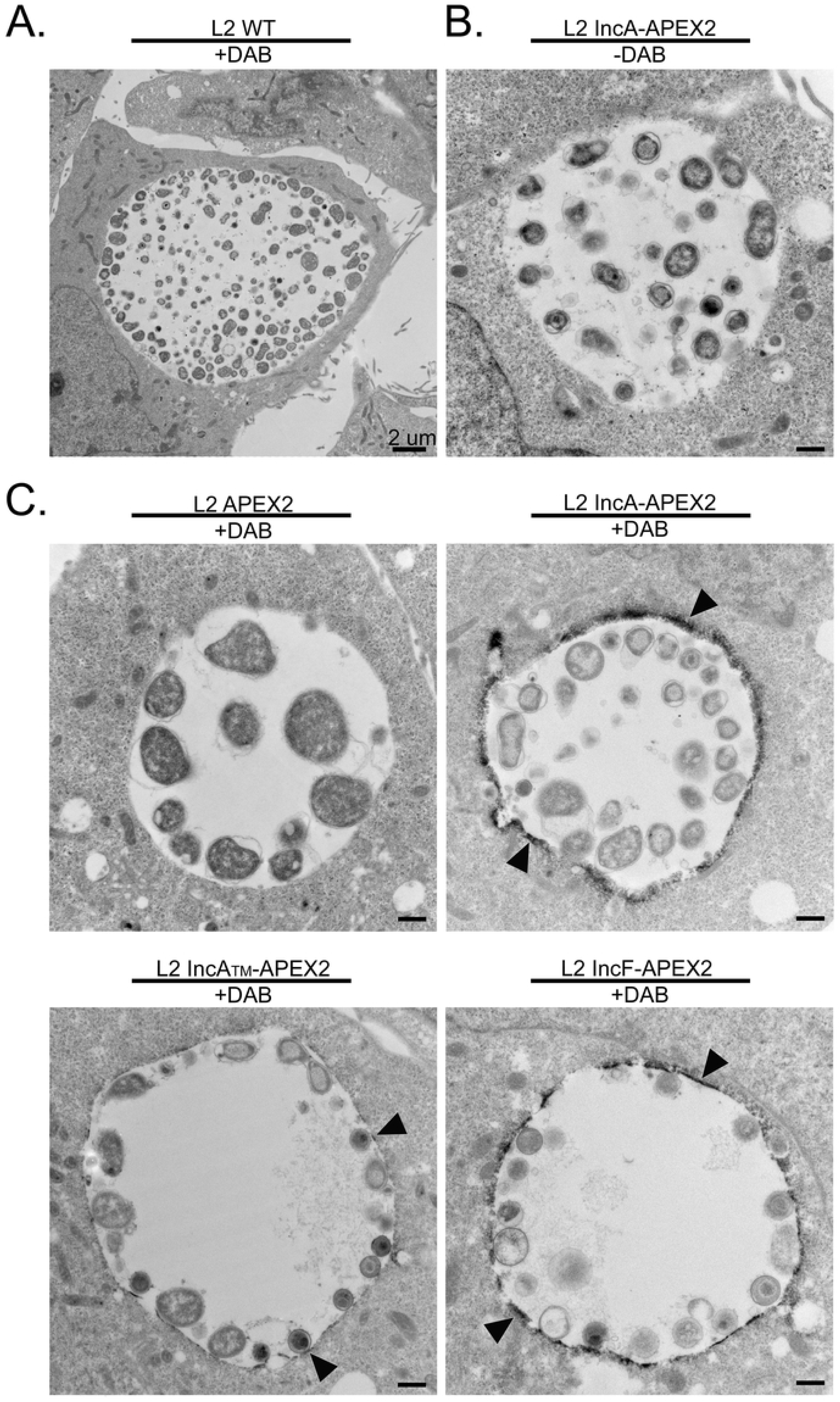
Ultrastructural localization of APEX2 activity to the cytosolic face of the inclusion membrane in HeLa cells infected with *C. trachomatis* L2 transformants expressing Inc-APEX2 constructs as determined by electron microscopy. HeLa cells seeded onto electron microscopy grade, cell culture treated coverslips were infected with *C. trachomatis* serovar L2 transformed with the indicated constructs, or *C. trachomatis* serovar L2 wild-type (WT) were induced with anhydrotetracycline (aTc) at 7 hpi (IncF-APEX2 0.3 nM aTc, all others 5 nM aTc). At 24 hpi, a glutaraldehyde and paraformaldehyde fixing solution was added to each sample and incubated on ice. Next, samples were pre-treated with DAB (or not, as indicated) 30 min prior to labeling by the addition of H_2_O_2_ solution (also containing DAB) to catalyze DAB polymerization. The reaction was quenched with glycine and processed for electron microscopy as indicated in *Methods*. (A) *C. trachomatis* L2 wild-type (WT) treated with DAB; scale bar= 2 µm, (B) *C. trachomatis* L2 IncA-APEX2 without DAB; scale bar= 500 nm, (C) *C. trachomatis* L2 transformants treated with DAB; DAB polymer staining around the inclusion is indicated by arrows; scale bar= 500 nm.

### Western blot detection of APEX2 containing constructs expressed from *C. trachomatis* L2 transformants

To confirm the correct expression of each construct containing APEX2, HeLa cells were infected with the *C. trachomatis* L2 APEX2, IncF-APEX2, IncA_TM_-APEX2, or IncA-APEX2 transformants and either not induced or induced at 7 hpi (0.3 nM aTc for IncF-APEX2 and 5 nM for all other transformants). Cell lysates were collected at 24 hpi and prepared for affinity purification using FLAG magnetic beads essentially as previously described (46). The eluates were blotted for the presence of each APEX2 containing construct using anti-FLAG antibody (the FLAG epitope tag is located at the N-terminus of APEX2). Lower levels of IncF-APEX2 (39.7 kDa) were detected compared to IncA_TM_-APEX2 (40.7 kDa), IncA-APEX2 (59.3 kDa), and APEX2 (30.3 kDa) (Fig S1). We detected some leaky expression of IncF-APEX2, IncA_TM_-APEX2, and IncA-APEX2 in our uninduced samples (Fig S1). As a loading control, the solubilized lysate was blotted for the presence of chlamydial Heat shock protein 60 (cHsp60) (Fig S1-Input, lower panel; cHsp60 antibody kind gift from Rick Morrison, University of Arkansas for Medical Sciences, Little Rock, AR). These data confirmed that each APEX2 containing construct was expressed from the *C. trachomatis* L2 transformants at the expected molecular weight.

### Affinity purification of biotinylated proteins

After confirming the correct construct localization, labeling activity at the inclusion membrane, and molecular weight of the proteins expressed from *C. trachomatis* L2, the lysates from *C. trachomatis* L2 IncF-APEX2, IncA_TM_-APEX2, IncA-APEX2 transformants, and the negative control infected HeLa cells from the biotinylation experiments described above (Fig 1) were affinity purified to isolate biotinylated proteins. The negative controls, mock-infected, *C. trachomatis* L2 wild-type infected, and *C. trachomatis* L2 APEX2 infected HeLa cells treated with biotin-phenol and hydrogen peroxide (to catalyze labeling) served to control for background, endogenous biotinylated proteins. As described previously, the major background endogenous biotinylated proteins include eukaryotic mitochondrial carboxylases (75 and 125 kDa) (30, 47), and, in *C. trachomatis* L2 infected HeLa cells, the chlamydial biotin ligase (21 kDa), which uses biotin as a co-factor (30, 48). We did not include uninduced *C. trachomatis* L2 transformants in our analysis because we observed some leaky construct expression and were concerned that using this as a negative control would subtract true interacting proteins during the analysis step (Fig S1).

Biotinylated proteins were affinity-purified using streptavidin beads and visualized by western blotting using a fluorescent streptavidin-conjugate (Fig 3; streptavidin panel). Biotinylated proteins were detected from each of the *C. trachomatis* L2 IncF-APEX2, IncA-APEX2, and IncA_TM_-APEX2 transformants that received both biotin-phenol and H_2_O_2_ (Fig 3; streptavidin panel). Without the addition of biotin-phenol to the *C. trachomatis* L2 Inc-APEX2 transformants, only endogenous biotinylated proteins were detected. Similarly, in each of the negative controls, *C. trachomatis* L2 APEX2, *C. trachomatis* L2 wild-type (not transformed) and mock-infected HeLa cells, only background endogenous biotinylated proteins were detected (Fig 3; streptavidin panel).

**Figure 3.**
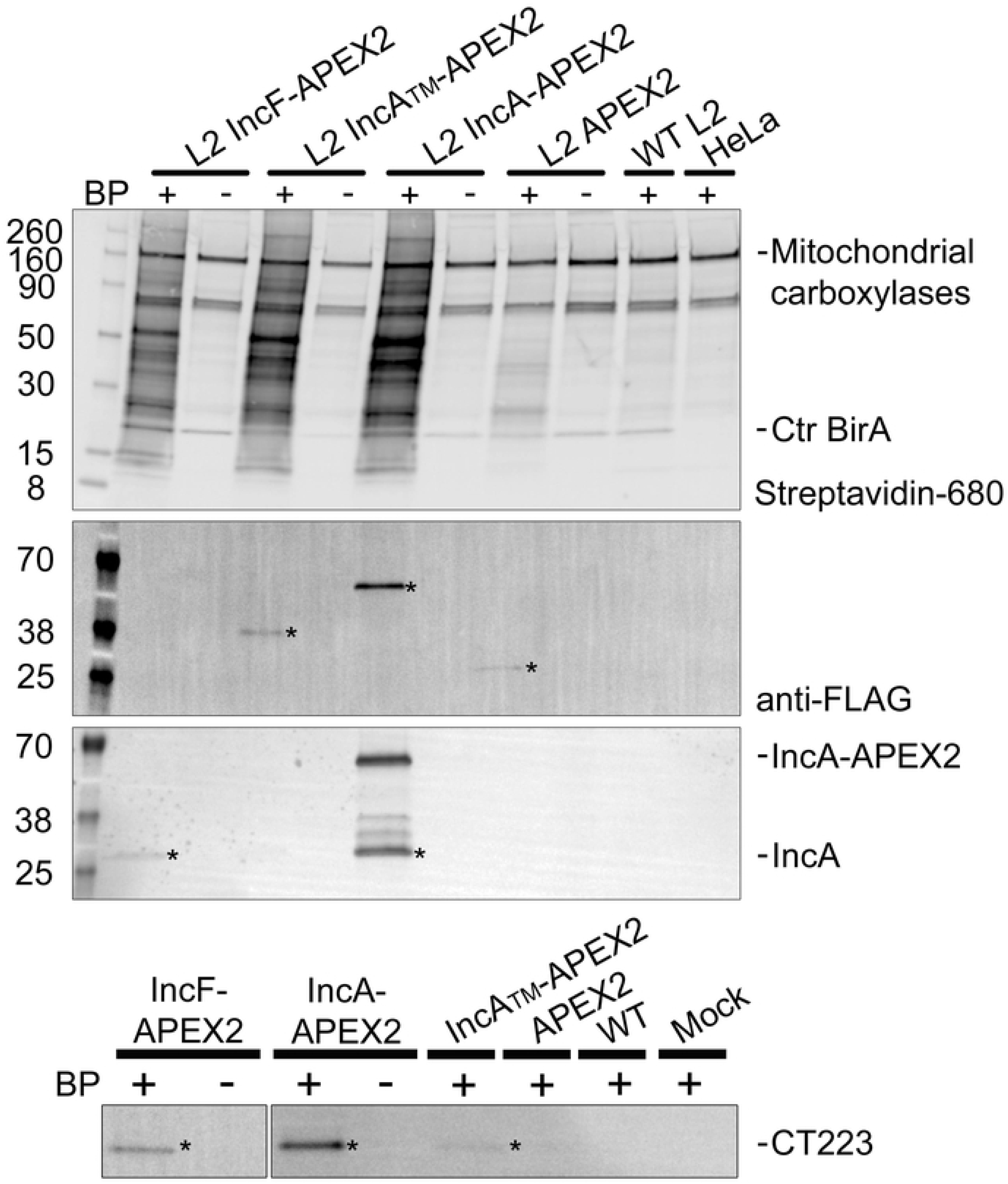
Western blot detection of affinity purified biotinylated proteins. HeLa cells infected with *C. trachomatis* L2 Inc-APEX2 transformants, wild-type (WT), or mock-infected were induced with anhydrotetracycline (aTc) at 7 hpi (0.3 nM for IncF-APEX2, and 4 nM for all others). Biotin-phenol (BP) was added 30 min prior to the biotinylation reaction at 24 hpi. Biotinylation was catalyzed by the addition of 3 mM H_2_O_2_ for 1 min and stopped with a quenching wash solution. Biotinylated proteins were affinity purified from solubilized lysates using streptavidin beads, eluted in sample buffer, separated by SDS-PAGE and transferred to PVDF for western blotting. The eluate fraction was probed for biotinylated proteins (streptavidin-680 conjugate), construct expression (anti-FLAG antibody), IncA (anti-IncA antibody), CT223 (anti-CT223 antibody), and imaged using Azure c600. Asterisks indicate detected proteins. See Supplementary Figure 1.

To determine if the expressed constructs containing APEX2 were biotinylated *in vivo* and affinity purified, we blotted the eluates using an anti-FLAG antibody (APEX2 contains the FLAG epitope in the N-terminus). We detected biotinylated IncA_TM_-APEX2 (40.7 kDa), IncA-APEX2 (59.3 kDa), and APEX2 (30.3 kDa) (Fig 3; FLAG panel). We did not observe biotinylated IncF-APEX2 (40.7 kDa) in the eluate fraction, which is likely a result of the lower expression necessary to preserve its correct localization ((30) and Fig S1). In addition, to determine if we could detect solubilized endogenous chlamydial Incs, we used an anti-IncA antibody (a gift from Dr. T. Hackstadt, NIAID, Rocky Mountain Laboratories, Hamilton, MT) and an anti-CT223 antibody (a gift from R. Suchland, University of Washington, WA and Dr. D. Rockey, Oregon State University, OR) to blot the eluates from the streptavidin affinity purification. We detected endogenous IncA in the eluate from *C. trachomatis* L2 IncF-APEX2 and IncA-APEX2 (Fig 3; IncA panel). The IncA antibody is specific for the C-terminus, so it detects IncA-APEX2 (59.3 kDa band) containing full-length IncA and not the truncated IncA_TM_-APEX2 expressed construct, which lacks the epitope that is recognized by the antibody. We also detected CT223 (29.3 kDa) in the streptavidin affinity purified eluate from each of the *C. trachomatis* L2 Inc-APEX2 samples but not in the negative controls (Fig 3C; CT223 panel). These western blot data provide an initial validation of our proximity labeling system because IncA homotypic interactions have been described (23, 37, 39, 40, 49) (e.g., IncA-APEX2 interacts with endogenous IncA in the inclusion membrane). These data are also consistent with previously published *in vivo* protein-protein interaction data using the BACTH system that identified IncF and IncA interactions (23).

### Mass spectrometry identification of streptavidin affinity purified biotinylated *C. trachomatis* L2 and eukaryotic proteins

To identify the proteins proximal to or interacting with the inclusion membrane that were biotinylated *in vivo* using the APEX2 proximity labeling system, the eluates from streptavidin affinity purification were briefly electrophoresed, sectioned, and then processed for mass spectrometry identification. To enhance mass spectrometry peptide identification, individual gel sections were digested with two enzymes, trypsin and AspN (50), and then processed as indicated in the *Methods*. Five biological replicates for each condition were analyzed by tandem mass spectrometry (MS/MS) and individual peptides were identified by performing Mascot searches against the *C. trachomatis* L2 (434/Bu) database and the *Homo sapiens* database. Our analysis detected 810 *C. trachomatis* L2 proteins (Table S1) and over 5,000 eukaryotic proteins (Table S2) in total from the combined datasets. To analyze our mass spectrometry data for statistical significance and to remove non-specific or background biotinylated proteins, we used Significance Analysis of INTeractome (SAINT) (38). SAINT uses quantitative data embedded in the raw mass spectrometry data from label-free quantification methods to filter out background peptides (38). The peptide spectra for a protein (i.e. prey) identified in the sample of interest (i.e. bait) is normalized to both the protein length and the total number of spectra compared to the negative controls. Bayesian statistics are used to calculate the probability of an interaction between each bait-prey interaction identified. The calculated probability is expressed as Bayesian False Discovery Rate (BFDR). We used BFDR less than or equal to 0.05 as a strict cut-off for our analysis parameters.

When we analyzed the *C. trachomatis* L2 proteins for statistical significance, several Inc proteins were among the top SAINT identified significant hits using our Inc-APEX2 constructs (Table 1; Table S1). Using our strict BFDR cut-off (BFDR≤0.05), we identified three statistically significant chlamydial proteins using *C. trachomatis* L2 IncF-APEX2 and IncA-APEX2, and two significant proteins using *C. trachomatis* L2 IncA_TM_-APEX2. CT223 was the only chlamydial protein that was identified as statistically significant using each *C. trachomatis* L2 Inc-APEX2 transformant. IncA was detected using *C. trachomatis* L2 IncA-APEX2 and IncA_TM_-APEX2. The identification of CT223 and IncA by mass spectrometry using IncA-APEX2 is supported by the detection of proteins eluted from the streptavidin affinity purified lysate (Fig 3). Statistically significant chlamydial proteins that were unique to the individual *C. trachomatis* L2 transformants included IncD and IncF which were identified using *C. trachomatis* L2 IncF-APEX2 and outer membrane complex B (OmcB) which was identified using *C. trachomatis* L2 IncA-APEX2 (Table 1; Table S1). Additional chlamydial Inc proteins that were detected by mass spectrometry but did not make the strict BFDR (BFDR≤0.05) cut-off using *C. trachomatis* L2 IncA-APEX2 include IncC (BFDR=0.09), CT813 (BFDR=0.1), IncD (BFDR=0.11), and IncE (BFDR=0.2) (Table S1). In contrast, there were no additional Incs identified using *C. trachomatis* L2 IncA_TM_-APEX2 with a less stringent cut-off (BFDR ≤0.2). Using *C. trachomatis* L2 IncF-APEX2, IncA (BFDR= 0.12), CT228 (BFDR= 0.15), and IncE (BFDR=0.18) were detected (Table S1). Although IncA was not statistically significant using IncF-APEX2, IncA was detected in the affinity-purified eluate of IncF-APEX2 by western blot (Fig 3; Table S1). These data are also supported by previously observed IncF-IncA interactions by BACTH (23) and IncA-IncA interactions that have been previously described (23, 39). Importantly, our AP-MS data analyzed against the *C. trachomatis* L2 (434/Bu) database were supported by our western blot data.

**Table 1.** Significant C. trachomatis L2 proteins

| Sample | Protein (Uniprot ID) | Gene name (Uniprot ID) | Protein name <sup>a</sup> | BFDR <sup>b</sup> |
| --- | --- | --- | --- | --- |
| **IncF-APEX2** | A0A0H3MKT3_CHLT2 | CTL0476 | CT223 | 0 |
|  | INCD_CHLT2 | incD CTL0370 | IncD | 0.02 |
|  | INCF_CHLT2 | incF CTL0372 | IncF | 0.03 |
| **IncA<sub>TM</sub>-APEX2** | A0A0H3MD02_CHLT2 | incA CTL0374 | IncA | 0 |
|  | A0A0H3MKT3_CHLT2 | CTL0476 | CT223 | 0.02 |
| **IncA-APEX2** | OMCB_CHLT2 | omcB omp2 CTL0702 | omcB | 0 |
|  | A0A0H3MKT3_CHLT2 | CTL0476 | CT223 | 0 |
|  | A0A0H3MD02_CHLT2 | incA CTL0374 | IncA | 0 |

<sup>a</sup> Protein name indicated using *C. trachomatis* serovar D naming convention
<sup>b</sup> SAINT Bayesian False Discovery Rate

When we applied SAINT to our *Homo sapiens* AP-MS data, 13 statistically significant eukaryotic proteins (BFDR ≤0.05) were identified using *C. trachomatis* L2 IncF-APEX2, 18 statistically significant proteins using IncA_TM_-APEX2, and 192 statistically significant proteins using IncA-APEX2 (Table S2 and S3). To visualize common pathways for eukaryotic protein biological processes and molecular functions, the significant eukaryotic proteins (BFDR ≤0.05) from each SAINT analyzed Inc-APEX2 dataset were evaluated by ClueGO (Cytoscape (51)) (Fig 4). For IncF-APEX2 (Fig 4A; Fig S2A) the 13 significant eukaryotic proteins identified were associated with transport and the negative regulation of biological processes. For IncA_TM_-APEX2, the 18 statistically significant proteins were associated with regulation of metabolic processes and biological processes (Fig 4B; Fig S2B). Finally, the pathway analysis of the 192 significant eukaryotic proteins for IncA-APEX2 yielded globally enriched pathways including regulation of cellular protein metabolic processes, vesicle-mediated transport, actin cytoskeleton organization, regulation of cellular component organization, and translation (Fig 4C; Fig S2C).

**Figure 4.**
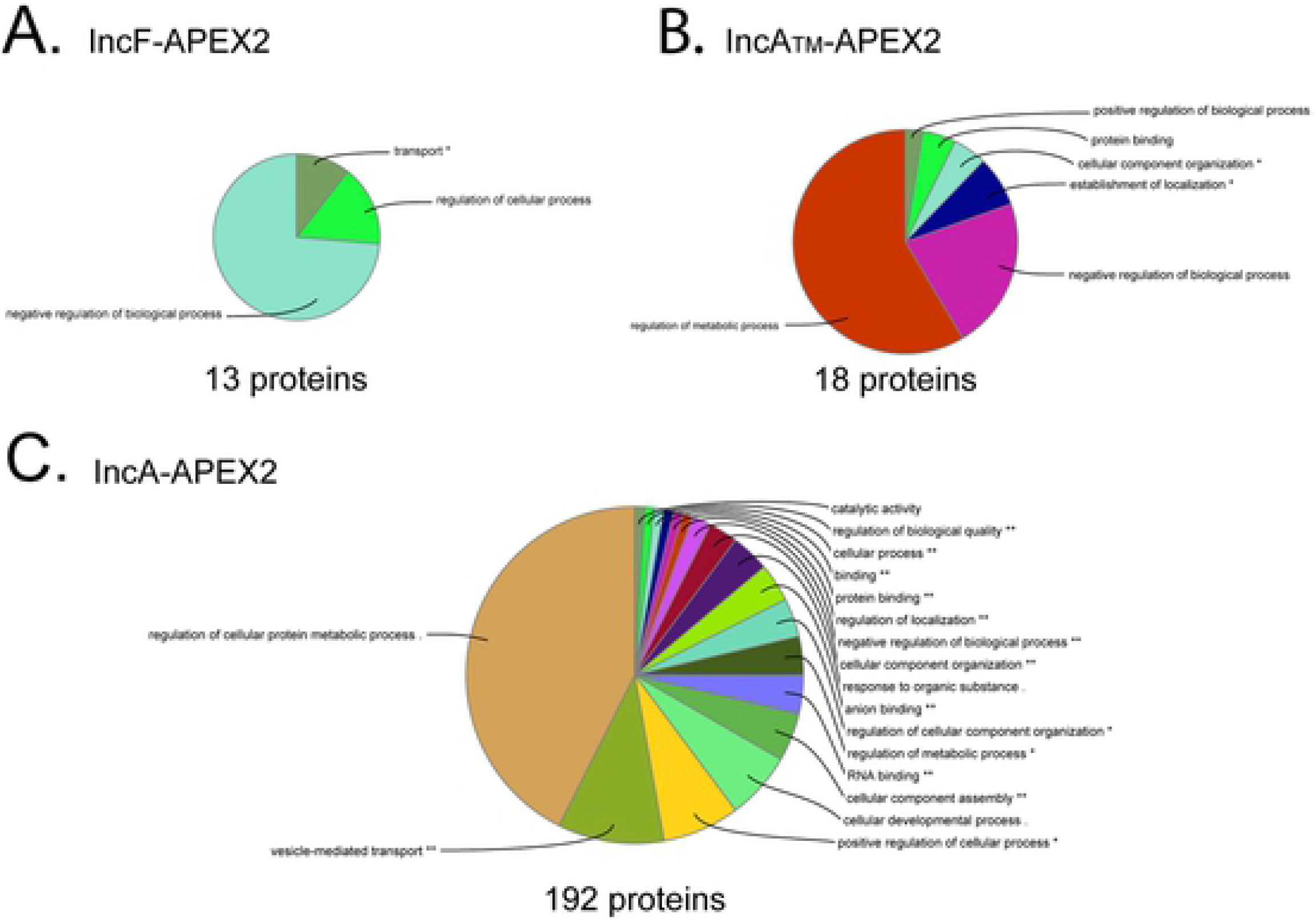
Visualization of global biological functions of AP-MS identified and statistically significant eukaryotic proteins from Inc-APEX2 pulldowns. ClueGO global network visualization of eukaryotic proteins identified by mass spectrometry (SAINT BFDR ≤ 0.05) from *C. trachomatis* L2 (A) IncF-APEX2, (B) IncA_TM_-APEX2, and (C) IncA-APEX2. See Supplementary Figure 2 and 3.

Individual datasets were also analyzed using STRING (Cytoscape (51)) to visualize the protein binding partner network for statistically significant (BFDR ≤0.05) eukaryotic proteins within each IncF-APEX2 (Fig S3A), IncA_TM_-APEX2 (Fig S3B), and IncA-APEX2 (Fig S3C) dataset. Four statistically significant eukaryotic proteins were common to all Inc-APEX2 datasets: Leucine-Rich Repeat Flightless-Interacting Protein 1 (LRRF1 or LRRFIP1), microtubule-associated protein 1B (MAP1B), Cystatin-B (CYTB), and brain acid soluble protein 1 (BASP1) (Table S2). Twelve proteins were shared between *C. trachomatis* L2 IncA-APEX2 and IncA_TM_-APEX2 including myosin phosphatase target subunit 1 (MYPT1 or PPP1R12A), transitional endoplasmic reticulum ATPase (TERA, VCP), microtubule-associated protein 4 (MAP4), multifunctional protein ADE2 (PUR6), sorting nexin-1 (SNX1), src substrate cortactin (SRC8, CTTN), methylosome protein 50 (MEP50), sorting nexin 6 (SNX6), perilipin-3 (PLIN3), eukaryotic translation initiation factor 4B (IF4B), nucleoside diphosphate kinase A (NDKA), and nucleoside diphosphate kinase B (NDKB) (Table S2). In both the IncF-APEX2 and IncA-APEX2 datasets, four eukaryotic proteins were statistically significant: 14-3-3-η (YWHAH), myristoylated alanine-rich C-kinase substrate (MARCKS), 14-3-3-β (YWHAB), and keratin type I cytoskeletal 20 (K1C20) (Table S2). These data included statistically significant eukaryotic proteins that have been previously shown to be recruited to the inclusion by Inc proteins. For example, in our IncA-APEX2 and IncF-APEX2 dataset, we identified 14-3-3-β, which is known to bind IncG (9). In addition, the eukaryotic proteins, SNX5 and SNX6, which bind IncE (24), and MYPT1, which binds CT228 (14, 15), were identified in both the IncA-APEX2 and IncA_TM_-APEX2 datasets (Table S2). Furthermore, the known chlamydial Inc binding partners for the eukaryotic proteins listed above (IncG, IncE, and CT228) were also identified in the AP-MS *C. trachomatis* L2 protein analyzed datasets (Table S1). We also identified eukaryotic proteins that are known to localize at the inclusion but for which an Inc binding partner has not been identified, including microtubule-associated protein 1B (MAP1B) (52) and Src-substrate cortactin (SRC8, CTTN) (53). Importantly, besides identifying eukaryotic proteins that are known to localize at the inclusion membrane, our Inc-APEX2 data identified several eukaryotic proteins that have not been previously examined for localization to the chlamydial inclusion.

### Co-localization of LRRF1 with the *C. trachomatis* L2 inclusion membrane

One of the high confidence AP-MS identified eukaryotic proteins (significant in each Inc-APEX2 dataset (BFDR=0)), Leucine-Rich Repeat in Flightless Interacting Protein 1 (LRRF1), has been reported to be involved in activating a type 1 interferon response (54-58), which plays a role in host cell clearance of intracellular bacteria during infection and the development of adaptive immunity (59). Also, a known LRRF1 binding partner called Protein Flightless-1 homolog (FLI1) (BFDR=0.02) (56, 60) was identified as significant by SAINT analysis in the IncA-APEX2 dataset (Table S2). FLI1 has been reported to associate with β-catenin to regulate its activity (61). In support, LRRF1 (24, 29, 33) and FLI1 (24, 29) were also identified in previous AP-MS and inclusion-MS experiments. Endogenous LRRF1 has not been examined for localization to the chlamydial inclusion.

LRRF1 was first confirmed by western blot (dimer 160 kDa) in the eluate from streptavidin affinity purified lysate from each of the *C. trachomatis* L2 Inc-APEX2 infected HeLa cells but not in the *C. trachomatis* L2 Inc-APEX2 samples that did not receive biotin-phenol, in the *C. trachomatis* L2 wild-type, or in mock-infected negative control samples (Fig 5A). These data confirm the identification of LRRF1 by mass spectrometry. To assess if LRRF1 and FLI1 localized to the chlamydial inclusion, HeLa cells were infected with *C. trachomatis* L2 wild-type, fixed at 24 hpi, and stained for immunofluorescence. LRRF1 (Fig 5B) and FLI1 (Fig 5C) were observed to localize to the inclusion membrane at 24 hpi. Subsequently, we transfected HeLa cells with a vector encoding LRRF1-GFP or FLI1-GFP and then infected or not with *C. trachomatis* L2 wild-type. At 24 hpi, cells were fixed and processed for immunofluorescence. In *C. trachomatis* L2 infected HeLa cells, LRRF1-GFP (Fig S4A) and FLI1-GFP (Fig S4B) were each observed at the inclusion membrane. In support, ectopically expressed HA epitope-tagged FLI1 was also reported at the *C. trachomatis* L2 inclusion (29). In mock-infected HeLa cells, LRRF1-GFP and FLI1-GFP both appeared diffusely in the host cytosol (Fig S4A and S4B, respectively). When we knocked down LRRF1 expression in HeLa cells, we did not observe a biologically significant change in the production of infectious progeny (Fig S4C; Non-targeting siRNA= 2.73 × 10^6^ IFU/mL; GAPDH siRNA=4.46 × 10^6^ IFU/mL; Single LRRF1 siRNA= 2.3 × 10^6^ IFU/mL; Pooled LRRF1 siRNA= 4.09 × 10^6^ IFU/mL).

**Figure 5.**
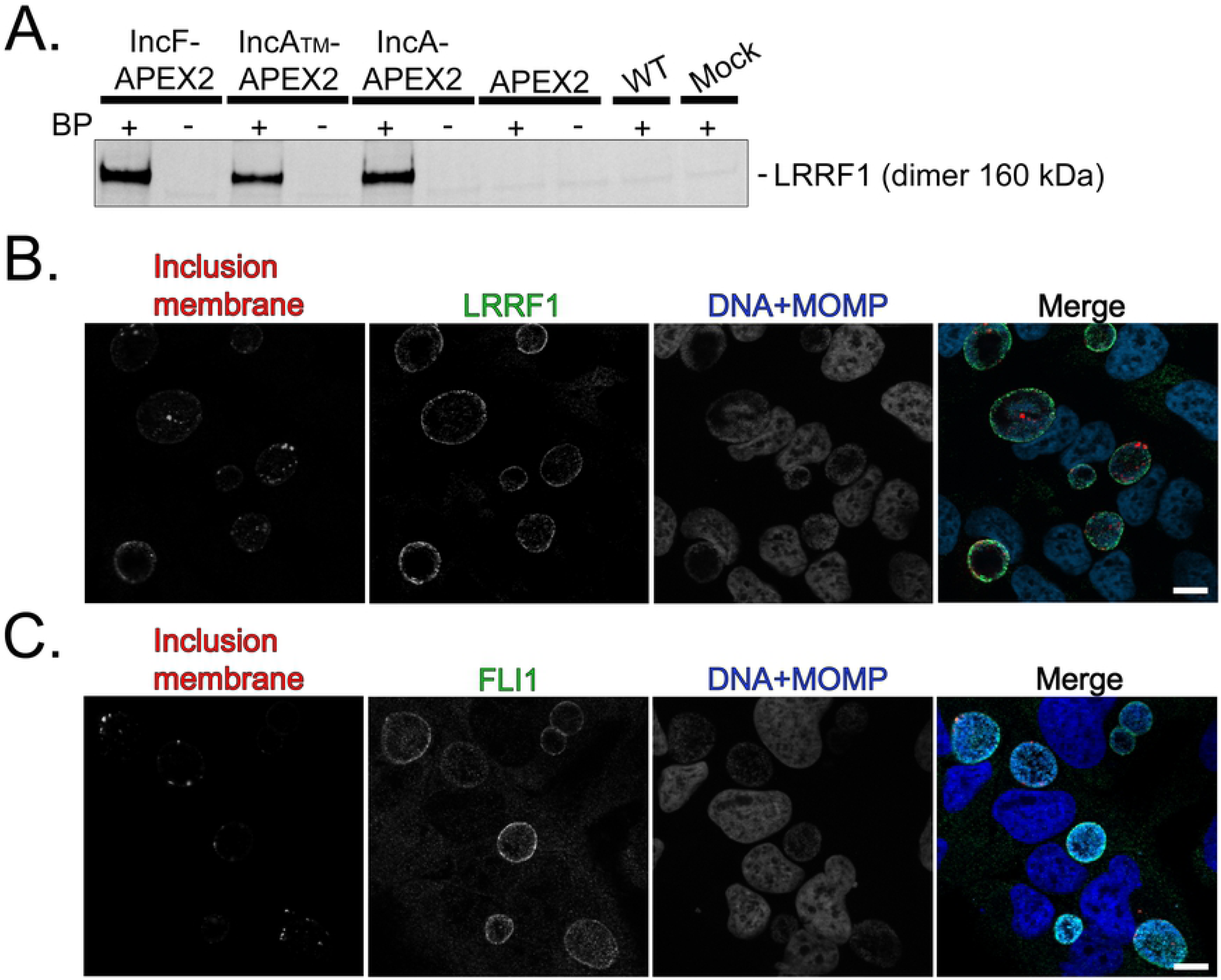
Confirmation of LRRF1 biotinylation by Inc-APEX2 proteins and localization of LRRF1 and FLI1 to the chlamydial inclusion. (A) Western blot confirmation of LRRF1 in the eluates from streptavidin affinity purified biotinylated lysate from *C. trachomatis* L2 IncF-APEX2, IncA_TM_-APEX2, and IncA-APEX2 transformants at 24 hpi (BP= biotin-phenol). (B) Confirmation of LRRF1 co-localization with the inclusion of *C. trachomatis* L2 wild-type infected HeLa cells. Cells were fixed at 24 hpi in 4% paraformaldehyde and permeabilized with 0.5% Triton X-100 then stained for indirect immunofluorescence to visualize the inclusion membrane (CT223; red), LRRF1 (green), DNA and *Chlamydiae* (DRAQ5 and MOMP; blue). (C) Confirmation of FLI1 co-localization with the inclusion of *C. trachomatis* L2 wild-type infected HeLa cells. Cells were fixed at 24 hpi in 4% paraformaldehyde and permeabilized with 0.5% Triton X-100 then stained for indirect immunofluorescence to visualize the inclusion membrane (CT223; red), FLI1 (green), and DNA and *Chlamydiae* (DAPI and MOMP; blue). Coverslips were imaged using Zeiss Apotome.2 with 100x magnification. Scale bar= 10 µm. See Supplementary Figure 4.

### LRRF1 co-localizes with the *C. trachomatis* inclusion from mid to late developmental cycle

To determine if LRRF1 stably or transiently localized to the inclusion during the developmental cycle, we infected HeLa cells with *C. trachomatis* L2 wild-type, fixed cells at intervals between 8 hpi and 36 hpi, and then stained for immunofluorescence to observe LRRF1 localization. Using CT223 as an inclusion membrane marker, LRRF1 could be observed at the inclusion as early as 12 hpi (Fig 6; arrows) and remained at the inclusion up to 36 hpi (Fig 6). Chloramphenicol (Cm) was added at 8 hpi to inhibit bacterial translation, and this treatment abolished the localization of LRRF1 to the inclusion (Fig 6; Cm treated panel), suggesting that LRRF1 recruitment is dependent on active chlamydial protein expression. These data indicate that LRRF1 is stably localized to the inclusion membrane from mid to late time points in the *C. trachomatis* L2 developmental cycle and that a chlamydial protein may recruit LRRF1 to the inclusion.

**Figure 6.**
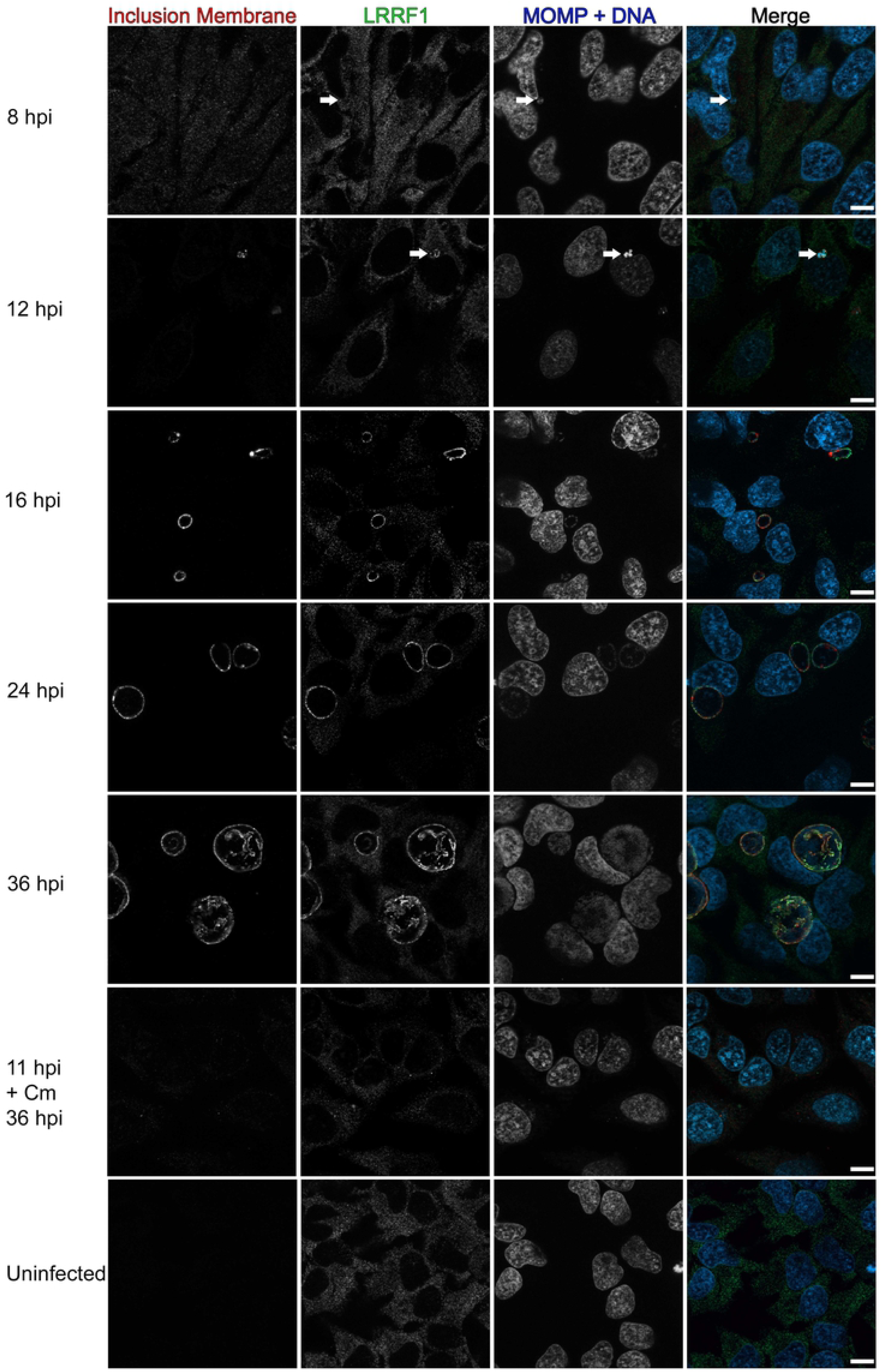
Recruitment of LRRF1 to the inclusion of *C. trachomatis* L2 during the developmental cycle and after chloramphenicol treatment. HeLa cells seeded on glass coverslips were infected with *C. trachomatis* L2 wild-type or mock infected. Wells were methanol fixed at 8, 12, 16, 24, and 36 hpi. One sample was treated with 34 µg/mL chloramphenicol (Cm) at 8 hpi and fixed at 36 hpi. Fixed coverslips were stained for indirect immunofluorescence to visualize LRRF1 (green), the inclusion membrane (CT223; red), and DNA and *Chlamydiae* (DAPI and MOMP; blue). Coverslips were imaged using a Zeiss ApoTome.2 with 100x magnification. Scale bar = 10 µm. Arrows indicate early inclusions at 8 hpi and LRRF1 co-localization with the inclusion at 12 hpi, respectively.

### LRRF1 co-localization with the inclusion is conserved among several *Chlamydia trachomatis* serovars and *Chlamydia* species

LRRF1 contains a coiled-coil domain as well as a cytosolic nucleic acid binding domain (55, 57), indicating two possible modes of LRRF1 recruitment to the inclusion membrane. To test if LRRF1 recruitment was mediated by a bacterial protein or as part of an innate response to infection by an intracellular bacterium, HeLa cells were infected, fixed, and processed as indicated in the *Methods* with various *Chlamydia trachomatis* serovars and *Chlamydia* species. The avirulent strain of *Coxiella burnetii* (Nine Mile Phase 2) was also included, which interacts with different eukaryotic pathways than *Chlamydia* (62). Our analysis of LRRF1 localization during infection of different *Chlamydia* species and *C. trachomatis* serovars revealed that LRRF1 co-localized with the inclusion of *C. trachomatis* serovar L2 (as observed above (Fig 6)), *C. trachomatis* serovar D, and the closely related *Chlamydia muridarum* (Fig 7A). LRRF1 did not co-localize with the inclusion of *Chlamydia pneumoniae*, *Chlamydia caviae*, or to the *Coxiella*-containing vacuole of *Coxiella burnetii* (Fig 7B). We conclude from these data that an Inc protein conserved between *C. trachomatis* serovar L2, serovar D, and *C. muridarum* recruits LRRF1 to the inclusion membrane.

**Figure 7.**
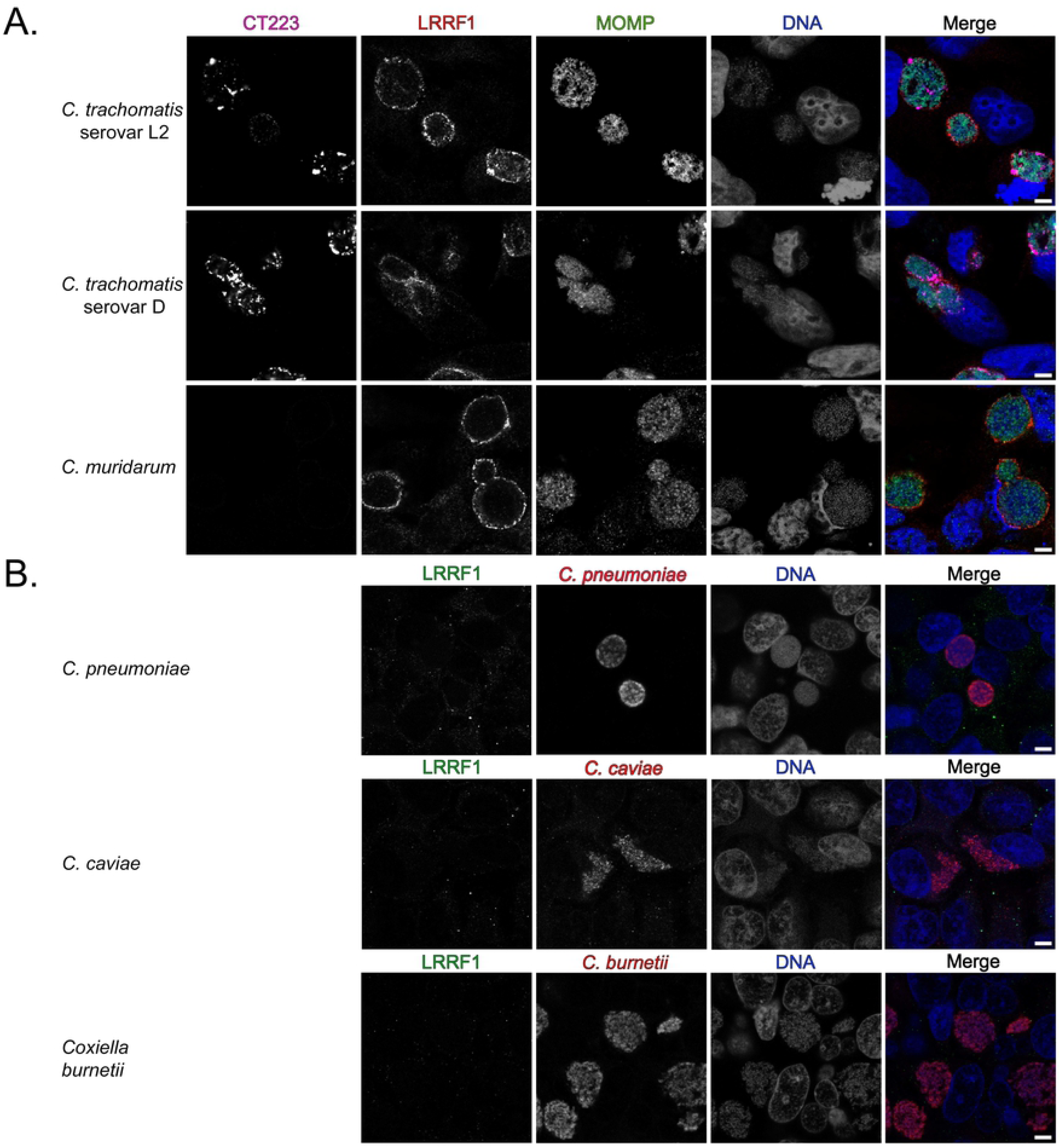
Examination of recruitment of LRRF1 to the inclusions of different chlamydial species and to the parasitophorous vacuole of the *Coxiella burnetii* Nine Mile Phase II. (A) HeLa cells were infected with *C. trachomatis* serovar L2, *C. trachomatis* serovar D, or *C. muridarum*, fixed with methanol at 24 hpi and stained for immunofluorescence to visualize the inclusion membrane (CT223; pink), LRRF1 (red), *Chlamydiae* (MOMP; green), and DNA (DAPI; blue). (B) HeLa cells were infected with *C. pneumoniae* and fixed in paraformaldehyde at 96 hpi, with *C. caviae* and methanol fixed at 24 hpi, or with *C. burnetii* Nine Mile Phase II, fixed with methanol at 3 days post-infection, and stained for immunofluorescence to visualize LRRF1 (green), bacteria (red), and DNA (DRAQ5; blue). Coverslips were imaged using a Zeiss confocal LSM 800 with 63x magnification and 2x zoom. Scale bar = 5 µm.

### BACTH assay to screen for LRRF1-Inc interacting partners

To determine if IncF and IncA used in the proximity labeling experiments can bind LRRF1, we used the bacterial adenylate cyclase two-hybrid (BACTH) system to screen for protein-protein interactions (23, 63, 64). Here, two plasmids encoding proteins of interest genetically fused to the catalytic fragments (i.e., T25 and T18) of the *Bordetella pertussis* adenylate cyclase are co-transformed into *E. coli* (Δ*cyaA*) (63, 65-67). An interaction between two proteins of interest brings the catalytic fragments in close proximity, restoring adenylate cyclase activity (63, 67, 68). Adenylate cyclase activity results in the production of cAMP and activates the expression of β-galactosidase via regulation of the chromosomally encoded *lac* operon in *E. coli* (63, 65-67). Positive interactions, indicated by the presence of blue colonies, are detected on minimal media (supplemented with Isopropyl β-D-1-thiogalactopyranoside (IPTG) and 5-Bromo-4-chloro-3-indolyl-β-D-galactopyranoside (X-gal), and interactions are quantified by β-galactosidase assay (63, 65-67).

A targeted screen was performed using IncF and IncA, CT223, CT813, CT288, and CT226. These Incs were either detected in our proximity labeling experiments or are Incs that are conserved between *C. trachomatis* serovar L2, serovar D, and *C. muridarum* (69). Of interest, LRRF1 contains a coiled-coil domain (57), which is a feature shared by several chlamydial Incs (39, 40, 70). Homotypic interactions have been previously described for IncA (23), which was used as a positive control. All interactions tested were quantified by β-galactosidase assay (23, 71). No interaction was observed between LRRF1 and IncF, IncA, CT288, CT223, or CT813 (Fig 8A). A positive interaction was detected between CT226 and LRRF1 (Fig 8A) and is consistent with previously reported data (24). CT226, like LRRF1, contains a coiled-coil domain (70). The interaction between LRRF1 and CT226 appeared specific because no other Incs tested, even Incs with coiled-coil domains, yielded a positive interaction (Fig 8A). In addition, CT226 and LRRF1 interactions were positive in both BACTH plasmid conformations (e.g., T25-LRRF1 vs. T18-CT226; T25-CT226 vs. T18-LRRF1).

**Figure 8.**
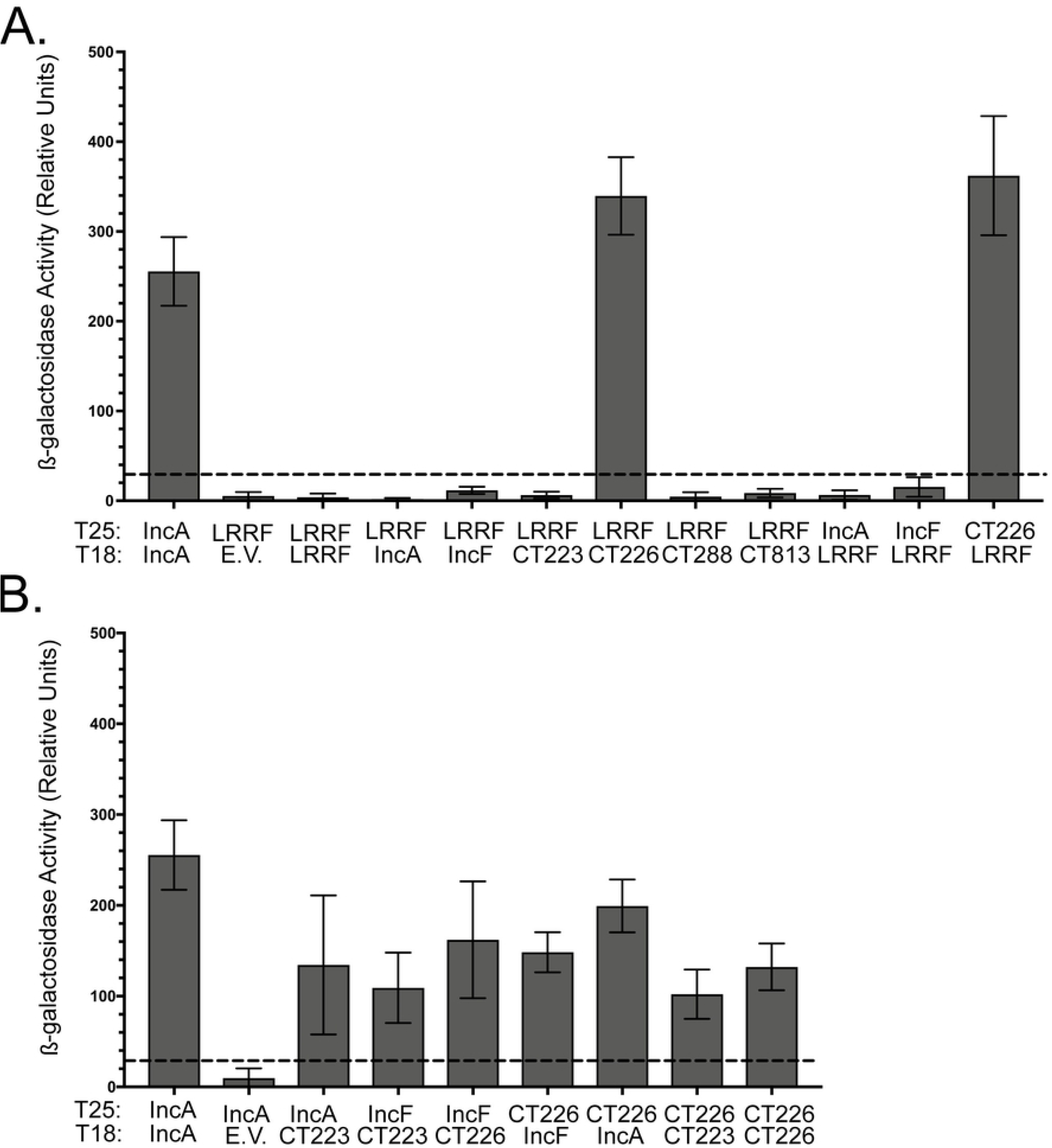
Bacterial Adenylate Cyclase Two-Hybrid (BACTH) assay to screen for LRRF1-Inc and Inc-Inc protein interactions. pST25 and pUT18 fused to chlamydial *Incs* or *LRRF1* as indicated were co-transformed into DHT1 *E. coli* (Δ *cyaA*), plated on minimal media containing IPTG and X-gal and grown for 3-5 days at 30 °C. Colonies were picked for overnight culture, and the interaction was quantified by β-galactosidase assay and reported at relative units (RU). (A) Quantitative analysis of LRRF1-Inc interactions, (B) Quantitative analysis of Inc-Inc interactions. Greater than five times the negative control is considered a positive interaction (indicated by the dotted line). Data shown are the mean and standard deviation from three biological replicates.

We did not detect a positive interaction between LRRF1 and IncA or LRRF1 and IncF, our original Inc-APEX2 constructs. Instead, it is possible that LRRF1 may be proximal to, but not directly binding, IncA and IncF at the inclusion membrane. To address this, we tested by BACTH the interactions of IncF and IncA with CT226, and both were found to interact with CT226 (Fig 8B). CT226 also demonstrated homotypic interactions (Fig 8B). Finally, we tested the ability of IncF and IncA to interact with CT223, the statistically significant Inc identified from each IncF-APEX2, IncA_TM_-APEX2, and IncA-APEX2 dataset. IncF and IncA each interacted with CT223 by BACTH (Fig 8B). CT223 also interacted with CT226 by BACTH (Fig 8B). These data support the possibility that CT223 and CT226 are proximal to IncF and IncA in the inclusion membrane. The identification of CT226 binding LRRF1 by BACTH assay corresponds to both the immunofluorescence data (Fig 7) and bioinformatic predictions because CT226 is conserved between *C. trachomatis* and *C. muridarum* but not *C. pneumoniae* or *C. caviae* (69).

### Assessment of LRRF1 co-localization with chlamydial Incs in *C. trachomatis* L2 infected HeLa cells by super-resolution microscopy

To assess the spatial localization and proximity of LRRF1 with respect to IncA and IncF, we used structured illumination (SIM) super-resolution microscopy. We also examined the localization of CT226, which was identified by BACTH as a potential interacting partner with LRRF1. Our IncA and IncF antibodies are both rabbit antibodies, as are the LRRF1 and FLI1 antibodies, precluding our ability to test endogenous IncF and IncA co-localization in *C. trachomatis* L2 infected eukaryotic cells. There are also no antibodies currently available to test if endogenous CT226 co-localizes with LRRF1 during *C. trachomatis* L2 infection. To assess co-localization of LRRF1 with IncF and IncA, we used our *C. trachomatis* L2 IncF-APEX2, IncA_TM_-APEX2, and IncA-APEX2 transformants. In addition, we created *C. trachomatis* L2 transformed with a plasmid encoding CT226 fused to a FLAG tag (CT226_FLAG_) to test localization between CT226 and endogenous LRRF1.

HeLa cells were infected with *C. trachomatis* L2 wild-type (i.e., non-transformed) or the transformants and induced for construct expression at 20 hpi (1 nM aTc for IncF-APEX2 and 5 nM for all other transformants). At 24 hpi, cells were fixed in ice cold methanol and processed for immunofluorescence as in *Methods* to detect the localization of IncF-APEX2, IncA_TM_-APEX2, IncA-APEX2, and CT226_FLAG_ with endogenous LRRF1. We assessed LRRF1 localization with endogenous CT223, the statistically significant chlamydial protein identified in each Inc-APEX2 dataset (Fig 9). Endogenous CT223 appeared in puncta as previously observed (72, 73), and LRRF1 appeared to uniformly localize around the inclusion, consistent with our earlier localization data for LRRF1 (Fig 5B). By SIM super-resolution microscopy, LRRF1 co-localized with each Inc-APEX2 construct, supporting the identification of LRRF1 using each construct (Fig 9). The expressed CT226_FLAG_ also co-localized with endogenous LRRF1 (Fig 9). Interestingly, the expression of CT226_FLAG_ resulted in fibers staining for CT226 extending from the inclusion, similar to what has been reported for IncA (74). LRRF1 was also observed to co-localize with CT226 fibers (Fig 9B; arrows indicate fibers). In contrast, LRRF1 did not co-localize with fibers from IncA-APEX2 transformants (Fig 9C).

**Figure 9.**
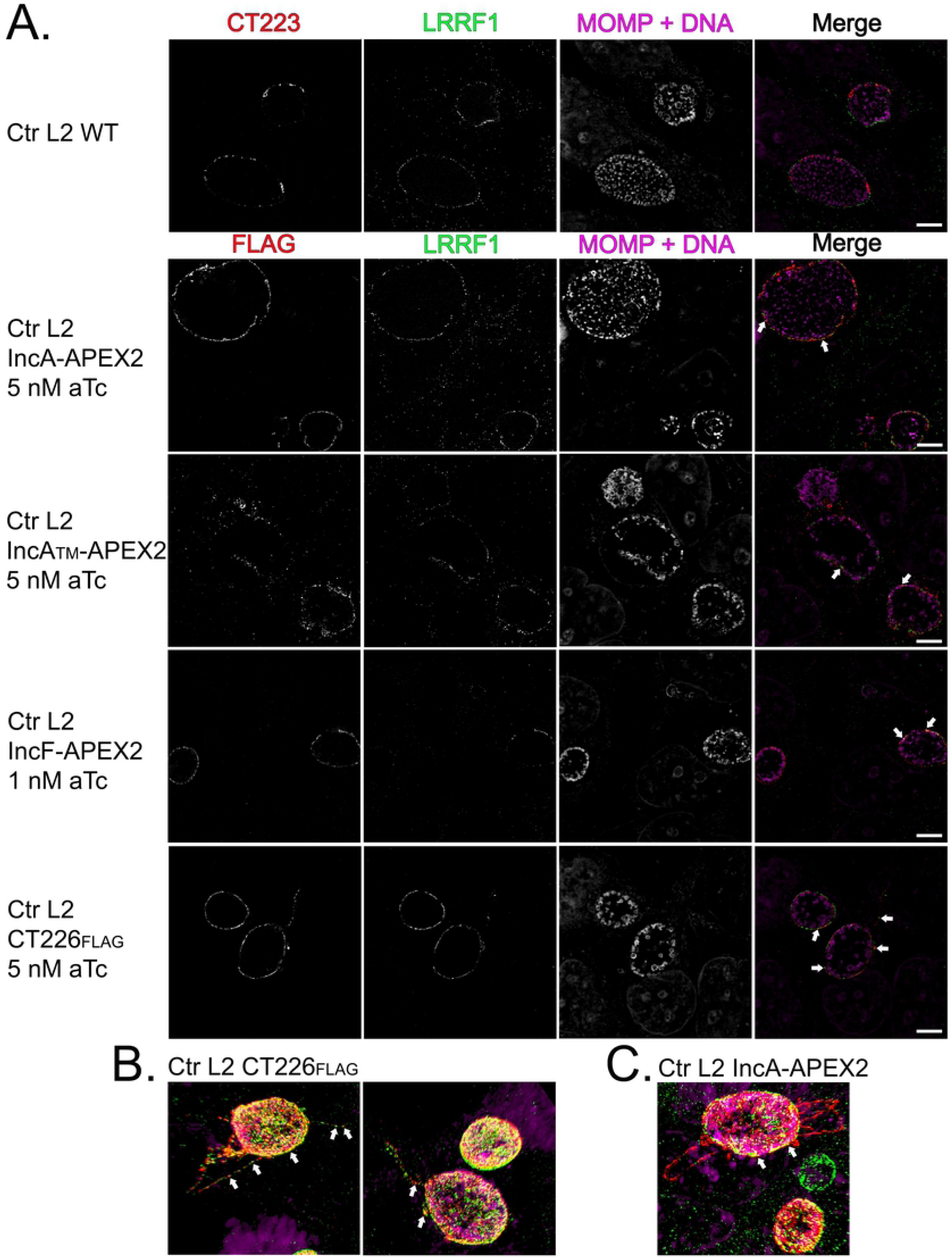
Assessment of LRRF1 co-localization with chlamydial Incs in *C. trachomatis* L2 infected HeLa cells using super-resolution microscopy. (A) Hela cells seeded on glass coverslips were infected with *C. trachomatis* L2 Inc-APEX2 transformants or CT226_FLAG_ transformants and induced for expression at 20 hpi (IncF-APEX2 was induced with 1 nM aTc; 5 nM aTc for all other transformants). At 24 hpi, coverslips were fixed with ice cold methanol and stained for immunofluorescence to visualize construct expression (FLAG) or CT223 (red), LRRF1 (green), *Chlamydiae* and DNA (DRAQ5 and MOMP; pink). Coverslips were imaged by Zeiss Elyra super-resolution microscopy 63×2x with structural illumination (SIM). Scale bar = 5 µm. (B) SIM 3D snapshot of *C. trachomatis* L2 CT226_FLAG_ infected HeLa cells with CT226_FLAG_ and LRRF1 positive fibers. (C) SIM 3D snapshot of *C. trachomatis* L2 IncA-APEX2 infected HeLa cells with IncA fibers. Arrows indicate co-localization between the indicated expressed construct and LRRF1.

### Co-immunoprecipitation of endogenous LRRF1 with *C. trachomatis* L2 CT226_FLAG_

To test if LRRF1 was directly binding to CT226, we performed co-immunoprecipitation assays with CT226_FLAG_ expressed from *C. trachomatis* infected HeLa cells. HeLa cells were plated in 6-well plates containing glass coverslips to confirm construct expression and localization. HeLa cells were infected with *C. trachomatis* L2 CT226_FLAG_ or IncF_FLAG_ transformants as a negative control. At 7 hpi, the constructs were either not induced or induced for expression using 5 nM aTc for *C. trachomatis* L2 CT226_FLAG_ and 1 nM aTc for IncF_FLAG_ (see (30) regarding IncF induction conditions). At 24 hpi, glass coverslips were removed, paraformaldehyde fixed, processed for immunofluorescence, and then cell lysates were collected and prepared for affinity purification using FLAG beads essentially as previously described (46). Both the clarified lysates (soluble fraction) and the eluates were blotted to detect each construct containing FLAG using an anti-FLAG antibody, and LRRF1 using an anti-LRRF1 antibody. Construct expression was observed for each *C. trachomatis* L2 CT226_FLAG_ or IncF_FLAG_ transformant by immunofluorescence co-localized with the inclusion membrane marker, IncA (Fig 10A). The FLAG affinity purified constructs were also detected by western blot (Fig 10B, CT226_FLAG_ 19.2 kDa, IncF_FLAG_ 11.3 kDa (monomer) and 22.6 kDa (dimer); Fig S5). However, LRRF1 (dimer 160 kDa) was only detected in the eluate fraction from the *C. trachomatis* L2 transformants induced for the expression of CT226_FLAG_ and not IncF_FLAG_. These data further support our BACTH data, indicating that LRRF1 can bind CT226_FLAG_ during *C. trachomatis* infection of eukaryotic cells.

**Figure 10.**
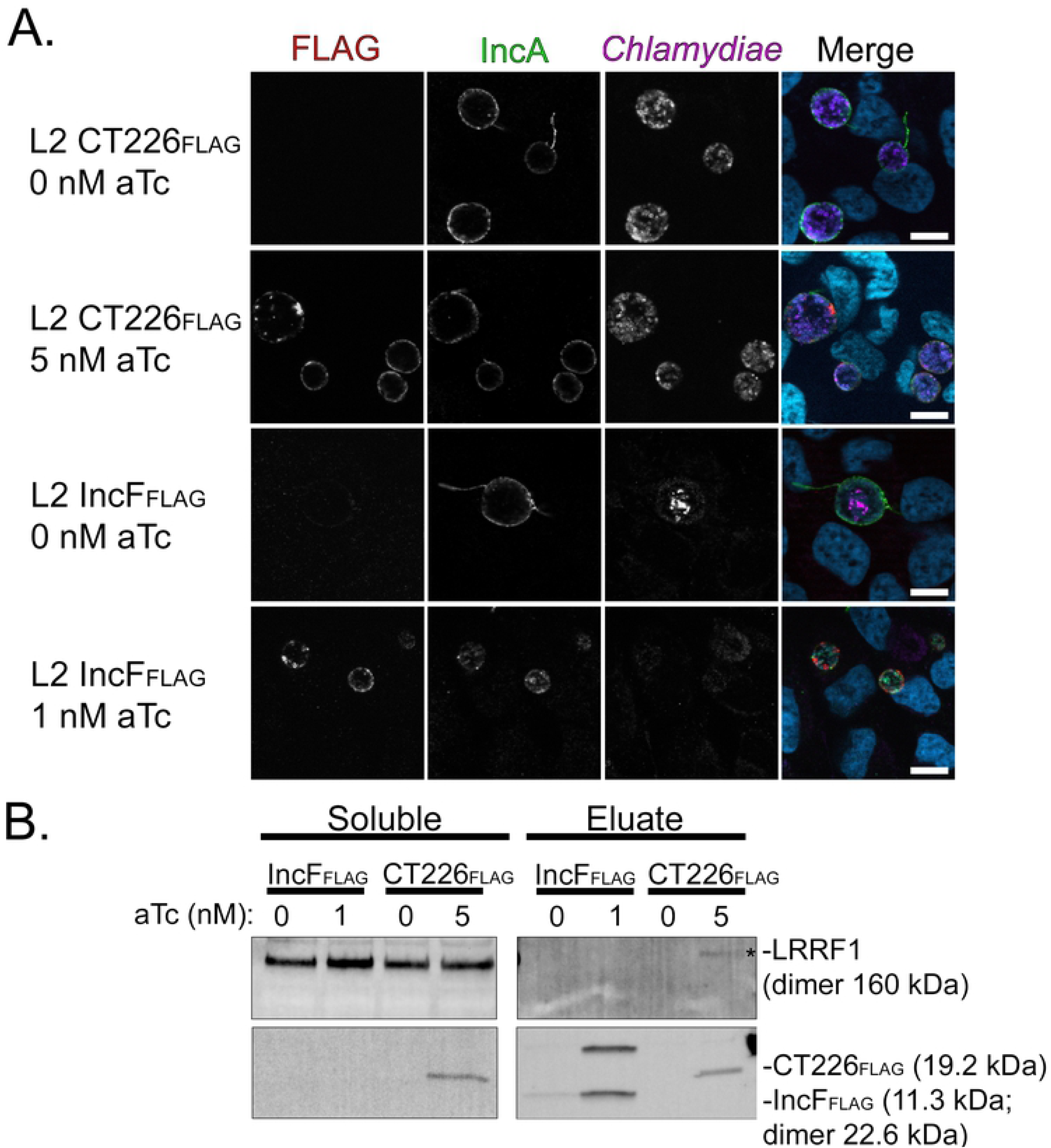
Co-immunoprecipitation of endogenous LRRF1 with *C. trachomatis* L2 expressed CT226_FLAG_. HeLa cells seeded in a 6-well plate with glass coverslips were infected with *C. trachomatis* L2 CT226_FLAG_ or IncF_FLAG_ and either not induced or induced for expression at 7 hpi with 5 nM aTc (CT226_FLAG_) or 1 nM aTc (IncF_FLAG_). (A) At 24 hpi, coverslips were removed, fixed in 4% paraformaldehyde, and stained to visualize FLAG (red), inclusion membrane marker (IncA; green), *Chlamydiae* (magenta), and DNA (blue). Coverslips were imaged using a Zeiss ApoTome.2 with 100x magnification. Scale bar = 10 µm. (B) The remaining cells were collected, solubilized, normalized, and affinity purified using FLAG beads. Clarified lysates (soluble) and eluates were probed for construct expression (FLAG; CT226_FLAG_ 19.2 kDa and IncF_FLAG_ 11.3 kDa), and LRRF1 (dimer 160 kDa). See supplementary figure 5.

## Discussion

We previously reported the feasibility of using the ascorbate peroxidase proximity labeling system (APEX2) in *C. trachomatis* L2 to detect protein-protein interactions at the inclusion *in vivo* (30). This tool improves upon past techniques to understand protein-protein interactions by maintaining the spatial organization of Inc proteins in the inclusion membrane (30). Proteins proximal to and within the inclusion membrane can be biotinylated and identified by affinity purification-mass spectrometry (AP-MS). Here, we used *C. trachomatis* L2 transformed with APEX2 fused to IncF and IncA, two Incs that, based on preliminary data, may represent distinct functional groups: Inc-Inc interactions to promote inclusion membrane organization and integrity or Inc-host protein interactions to facilitate chlamydial-host interactions and nutrient acquisition. As a control, we also prepared an IncA_TM_-APEX2 transformant that lacks the C-terminal SNARE-like domain of IncA and more closely resembles IncF in size.

As a field, we are at the early stages of understanding how the expression of various Inc constructs in the inclusion membrane can alter inclusion membrane organization and host-protein recruitment. We focused on expressing our Inc-APEX2 constructs under conditions similar to endogenous expression levels. This is an important consideration as overexpression of certain Incs can have deleterious effects on inclusion development and Inc localization (30) or can recruit a greater abundance of eukaryotic proteins (Fig 9B) that may or may not reflect *in vivo* conditions. This in contrast to a recent study where the authors over-expressed IncB-APEX2 (33), which did not result in localization consistent with endogenous IncB (28, 53). The goal of this study may have been to identify all possible inclusion interactions, regardless of specificity. A high confidence hit (see caveat noted below concerning statistical tools used) with their analysis conditions was Sec31A, yet the colocalization of this protein to inclusions with wild-type (untransformed) *C. trachomatis* was not strong (see Figure 6 of Dickinson et al., 2019 (33)). This does not necessarily diminish the findings of this group because the underlying reasons for this apparent discrepancy may reveal an important understanding of how the inclusion membrane and surrounding structures are organized. Here, we have sought to understand the context for why a specific protein was prominent in ours and others’ datasets, which ultimately revealed important information for how the inclusion membrane may be organized (Fig 11).

**Figure 11.**
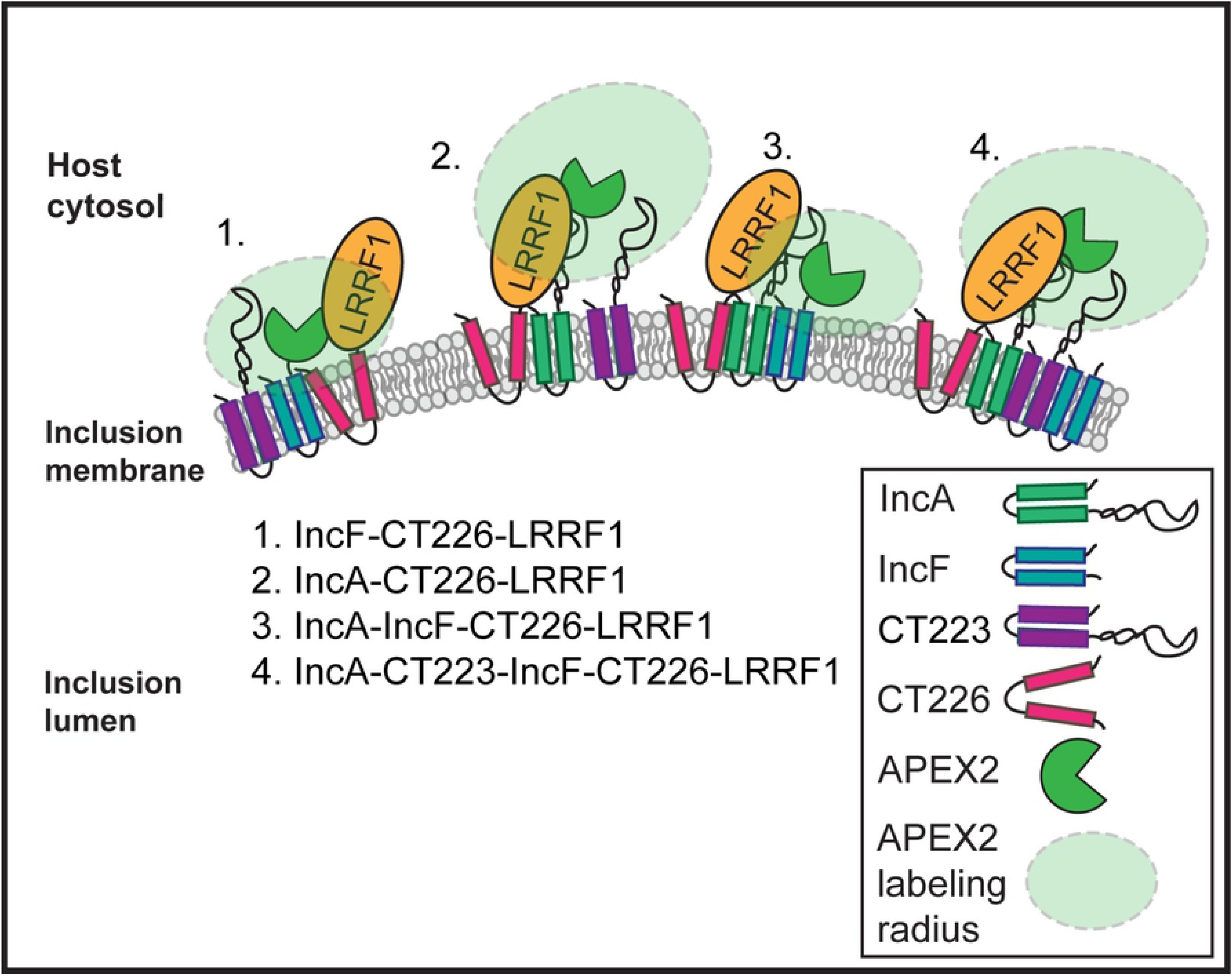
Model of Inc-Inc organization in the inclusion membrane and Inc-APEX2 proximity labeling. Proposed model of Inc organization based on mass spectrometry identified chlamydial Inc proteins using IncA-APEX2 and IncF-APEX2 proximity labeling constructs and bacterial two-hybrid assays (BACTH) to test protein-protein interactions. Based on these data we propose four possible scenarios for the spatial organization of Incs and how these Incs were detected using the APEX2 proximity labeling system: (1) IncF interacts with CT226 which binds LRRF1. (2) IncA interacts with CT226 which binds LRRF1. (3) IncA binds IncF and CT226 which binds LRRF1. (4) IncA, CT223, IncF, and CT226 (which binds LRRF1) all interact with each other. CT223 was statistically significant by SAINT analysis from mass spectrometry data and was able to interact with IncF and IncA by BACTH.

To assign statistical significance and eliminate background contaminant proteins from the AP-MS data in an unbiased fashion, the identified proteins from *H. sapiens* and *C. trachomatis* L2 were analyzed by Significance Analysis of INTeractome (SAINT) (38) (Table 1; Table S1-S3). This type of analysis is an improvement over the previously described statistical analyses used for similar datasets (33) because the t-test and G-test to determine whether to include a protein in the dataset is not sufficient to estimate the False Discovery Rate (FDR). Newer methods, such as PepC, use a matrix of t-test and G-test confidence intervals to detect differentially expressed proteins (75). The SAINT method constructs separate distributions for true and false interactions to derive the probability of the observed bait-prey interaction. The probability model for the bait-prey interaction pair is used to estimate measurement errors in a transparent manner. SAINT generates a Bayesian FDR (BFDR) calculation for each potential interaction detected in the dataset. SAINT also normalizes spectral counts based on protein length, which affect the potential availability of peptides that can be analyzed by MS/MS. These are more rigorous statistical tests than that obtained by independent t-test and G-test analyses (38, 76, 77).

We applied STRING interaction and ClueGo pathway analysis tools to our SAINT significant (BFDR≤0.05) eukaryotic proteins identified using the Inc-APEX2 constructs to detect globally enriched pathways. Consistent with our original hypothesis that IncA may preferentially interact with eukaryotic proteins compared to IncF, we detected a larger number of statistically significant eukaryotic proteins with our IncA-APEX2 (192 total) construct than with IncF-APEX2 (13 total), albeit with the caveats related to labeling radius noted below. For IncA_TM_-APEX2, there was more than a log reduction in the number of proteins identified when compared to full-length IncA-APEX2, suggesting specificity for interactions at the C-terminus of IncA. Given the presence of a SNARE-like domain in the C-terminus of IncA, it is possible that the large number of proteins identified with IncA-APEX2 reflects its interactions with other SNARE proteins on vesicles carrying diverse cargo. For instance, vesicle-mediated transport (e.g., ANXA1, AP1M1, CAV1, GOLGA2, PDCD6, PDCD6IP, RAB34, RAB5B, SEC16A, SEC24C, SEC31A, SNX1, SNX2, SNX3, SNX5, SNX6, TFG, TSG101, USO1) has been described in the context of *C. trachomatis* L2 acquisition of specific lipids from Golgi apparatus-derived exocytic vesicles (7, 10, 11). Statistically significant hits involved in SNX-retromer pathway disruption during chlamydial infection include SNX1, SNX2, SNX3, SNX5, SNX6, and SNX27 (24). Additional globally enriched biological processes and molecular functions involving cytoskeleton organization and translation align with previously published data. Cytoskeleton organization (e.g., ACTN1, ACTN4, CDC42, DPYSL3, DYNLL1, MARCKS, PLS3, RAC1, RHOA, SHTN1) corresponds with the literature as the inclusion is surrounded by an F-actin cage (78-80). One of the four statistically significant eukaryotic proteins identified using each Inc-APEX2 construct, microtubule-associated protein 1B (MAP1B) (Table S2) would be expected because microtubules are known to surround the inclusion (52, 81, 82) and thus would be proximal to both IncF and IncA, which uniformly label the inclusion (21). This interpretation is consistent with the findings of another study using APEX2 (33).

Our AP-MS data also identified multiple statistically significant Inc proteins. Again, IncF-APEX2 labeled three different Incs compared to only two for each of the IncA-APEX2 constructs (one of which was IncA itself). Consistent with our hypothesis, these data, taken together with the few eukaryotic proteins identified, suggest IncF may preferentially interact with other Inc proteins. Importantly, IncA_TM_-APEX2 did not label more Incs than full-length IncA, even with a less stringent BFDR threshold. These data indicate specificity to the IncF-APEX2 identified Incs. Although we identified chlamydial Incs with our Inc-APEX2 proximity labeling system, the majority of Incs detected were not statistically significant by SAINT (Table 1; Table S1). Some of this may reflect the residues that APEX2 covalently modifies during the biotinylation reaction (cysteine, histidine, tryptophan, and tyrosine residues) (31, 32, 45). The Incs that were significant (e.g., CT223 and IncA) have 11-20 cytosolically exposed target amino acids, whereas Incs that were not found to be statistically significant have fewer than 5-6 exposed target amino acids, in general. Therefore, proteins containing fewer APEX2-modifiable amino acids will not be efficiently tagged with biotin and subsequently not enriched as efficiently in the streptavidin affinity purification. Secondly, there are also inherent difficulties in identifying hydrophobic proteins by mass spectrometry. To counter this difficulty, we included two enzymes to digest purified proteins into peptides, but these efforts, in combination with limited modifiable amino acids, may not have allowed for enough enrichment of APEX2-targeted Inc proteins. To compensate for this limitation, a lower BFDR significance threshold could be considered when analyzing chlamydial Inc proteins. For example, when we lowered the BFDR threshold to 0.2, we identified eight Incs from our AP-MS data, including IncA in the IncF-APEX2 dataset that we detected by western blot (Fig 3). Further experimentation and analysis of Inc-Inc APEX2 data are required to identify an appropriate cut-off.

Using chlamydial expressed Inc-APEX2 constructs, we identified several chlamydial Incs and their known interacting eukaryotic protein partners including IncG and 14-3-3β (9), IncD and CERT (12, 13), CT228 and myosin phosphatase target protein subunit 1 (MYPT1) (14), and IncE and sorting nexin 5 (SNX5) and SNX6 (24) (Table 1; Table S1-S3). In addition, we identified eukaryotic proteins that were unique to each IncF-APEX2 and IncA-APEX2 datasets that have not been demonstrated to localize to the chlamydial inclusion, and, thus, require further validation (Table 2). In support of our hypothesis that IncA might preferentially interact with eukaryotic proteins compared to IncF, more eukaryotic proteins were identified using IncA-APEX2 than with IncF-APEX2 as noted above (Table S2). We expanded on the current knowledge of proteins recruited to the inclusion membrane with the validation of two eukaryotic proteins not previously reported at the inclusion: LRRF1, which was statistically significant in each Inc-APEX2 dataset, and its known binding partner FLI1, which was statistically significant in the IncA-APEX2 dataset (Table S2; Fig 5-7). We also identified a potential chlamydial protein interacting partner for LRRF1 by BACTH: CT226 (Fig 8A). These data are consistent with a previous report of LRRF1 and CT226 potential interactions identified by transfecting host cells with epitope-tagged CT226 followed by AP-MS (24).

We detected LRRF1 at the inclusion membrane, but LRRF1 knockdown does not negatively impact chlamydial progeny production in HeLa cells (Fig S4C). One explanation for this may be that *C. trachomatis* already prevents the normal function of LRRF1 by sequestering it at the inclusion membrane. In this context, knockdown would not affect the production of infectious progeny. Alternatively, LRRF1 has been implicated in the production of a type 1 interferon response (55, 56) so a more relevant tissue culture model may be required, such as human macrophages (83), which produce a robust interferon-mediated immune response. As no phenotype for LRRF1 knockdown was apparent, we chose to examine the nature of how LRRF1 was biotinylated by our constructs and ultimately identified in our dataset, as it has also been found in previous AP-MS datasets (24, 29, 33). The BACTH assays (Fig 8A) provide evidence for an interaction between LRRF1 and CT226. Further, the SIM super-resolution data indicated co-localization between LRRF1 and overexpressed CT226_FLAG_ (*C. trachomatis* L2 transformant) at the inclusion membrane as well as with CT226_FLAG_ positive fibers emanating from the inclusion (Fig 9). Lastly, we identified LRRF1 in CT226_FLAG_ co-immunoprecipitations, indicating that these proteins are true binding partners during chlamydial infection. We conclude that LRRF1 was likely labeled by APEX2 because CT226 is adjacent to, and likely interacting with, IncA and IncF in the inclusion membrane, which would position it within the labeling radius of our APEX2 constructs (Fig 11).

Our study has also revealed potential limitations of the APEX2 proximity labeling system to distinguish differences between Inc-protein binding partners and proteins that are in spatial proximity to the Incs at the inclusion membrane, as summarized in Figure 11. For instance, the labeling radius of APEX2, at least in our hands, may be larger than originally described (45). This would explain the identification of proteins proximal to IncF and IncA and not only specific protein binding partners at the inclusion (Fig 11). For example, although we identified strong LRRF1 recruitment to the inclusion (Fig 5), IncF and IncA, our bait proteins, were not identified as interacting partners of LRRF1 by BACTH or using IncF_FLAG_ co-immunoprecipitations (Fig 8A). In addition, the spatial organization of Incs in the inclusion membrane is not currently well understood, but IncA and IncF uniformly decorate the inclusion membrane as opposed to CT223, which localizes in discrete regions. *In vivo* two-hybrid experiments have shown that IncA and IncF interact (23), which might support the labeling of similar proximal proteins, and we identified IncA in IncF-APEX2 labeled eluates (Fig 3). Another possible explanation for a larger labeling radius is related to the diffusion rate and half-life of biotin-phenol, which is approximately one millisecond (32). Diffusion of biotin-phenol would contribute to a greater labeling radius and a larger pool of proteins identified. In support of the diffusion of biotin-phenoxyl radicals with our Inc-APEX2 constructs, we identified Outer Membrane Complex B (OmcB) and Major Outer Membrane Protein (MOMP) in the AP-MS data (Table 1, Table S1). It is possible that during labeling with Inc-APEX2 transformants, biotin-phenoxyl diffuses across the inclusion membrane and labels the bacteria (intra-inclusion) before the quenching step, and OmcB and MOMP are amongst the most abundant outer membrane chlamydial proteins. Shorter labeling times may decrease the labeling radius and increase labeling specificity (84). Also, BACTH assays are a useful tool to determine protein-protein interactions *in vivo* using *E. coli*. In this study, using the BACTH assay to test chlamydial Inc-Inc protein interactions, we observed CT223-IncF and CT223-IncA interactions (Fig 8B), which supports the identification of CT223 as a statistically significant Inc using each Inc-APEX2 construct (Table 1). However, we might miss some interactions with the BACTH system if eukaryotic post-translational modifications are required for the protein interactions to occur (e.g., phosphorylation; (85)). Therefore, other validation methods such as super-resolution microscopy or Duolink® PLA technology, provided antibodies are available, are required. Alternatively, overexpression models could be used to detect interactions with epitope-tagged Incs (with the caveats previously noted). Importantly, these data highlight the necessity of using adequate controls and statistical analyses to eliminate false positives and other proteins that may be transiently near the inclusion during the labeling period.

Our data highlight the utility of the ascorbate peroxidase proximity labeling system to detect novel protein interactions at the *C. trachomatis* inclusion membrane *in vivo*. This tool improves upon past techniques by maintaining the spatial organization of Incs in the inclusion membrane and biotinylating proximal proteins *in vivo*. Our goal was to determine if there is a preference for certain Incs toward Inc-Inc interactions or Inc-eukaryotic interactions in the inclusion membrane using the AP-MS SAINT data as the foundation for further study. Determining the complex types of interactions that Incs orchestrate in the inclusion membrane will lead to a better understanding of how *Chlamydiae* survive in their intracellular niche. Importantly, this technique is broadly applicable, when properly controlled, to other intracellular bacteria or parasites residing within a membrane-bound vacuole.

## Methods

### Antibodies and reagents

Primary antibodies used: mouse anti-FLAG (Sigma), rabbit anti-FLAG (Sigma), mouse anti-Giantin (Enzo), mouse anti-GAPDH (EMD Millipore), goat anti-MOMP (Meridian), rabbit anti-LRRF1 (Bethyl), rabbit anti-FLI1 (Thermo Fisher), rabbit anti-IncA (Kind gift from Ted Hackstadt, NIAID, Rocky Mountain Laboratories, Hamilton, MT), mouse anti-CT223 (Kind gift from R. Suchland, University of Washington, WA;, D. Rockey, Oregon State University, OR), mouse anti-*C. trachomatis* Hsp60 (a kind gift from Rick Morrison, University of Arkansas for Medical Sciences, Little Rock, AR), rabbit anti-*C. burnetii* (Elizabeth A. Rucks), mouse anti-*C. pneumoniae* AR39 (a kind gift from H. Caldwell, NIAID, Bethesda, MD). Secondary antibodies used for immunofluorescence were each donkey anti-647, 594, 488, and 405, or streptavidin-488 conjugate. DRAQ5 and DAPI were used to visualize DNA as indicated. Western blots were visualized using the appropriate secondary antibodies conjugated to IRDye 680LT, or IRDye 800 CW (LiCor Biosciences, Lincoln, NE), and membranes were imaged using Azure c600 (Azure Biosystems, Dublin, CA) and processed using Adobe photoshop creative cloud (Adobe).

### Organisms and cell culture

HeLa 229 cells [American Type Culture Collection (ATCC); Manassas, VA; CCL-2.1] were cultured at 37°C with 5% CO_2_ in biotin-free DMEM (Gibco; Grand Island, NY) that was supplemented with 10% heat-inactivated fetal bovine serum (FBS; HyClone, Logan, UT), for routine tissue culture, or with 1% FBS, for experiments involving biotinylation as previously described (30) with 10 μg/mL gentamicin (Gibco-BRL/Life Technologies; Grand Island, NY). HeLa cells were used to propagate *Chlamydia trachomatis* serovar L2 (LGV 434) for purification using established protocols (86, 87). Chlamydial titers were determined using conventional protocols to establish multiplicities of infection (m.o.i.), based on inclusion forming units (i.f.u.) and determined in HeLa cells as previously described (87, 88). McCoy cells (ATCC; Manassas, VA; CRL-1696) were cultured at 37°C with 5% CO_2_ in biotin-free DMEM (Gibco; Grand Island, NY) that was supplemented with 10% fetal bovine serum (FBS; HyClone, Logan, UT) used for *C. trachomatis* L2 (LGV 434) transformation experiments. HeLa cells, McCoy cells, and density-gradient purified *C. trachomatis* strains are routinely tested for *Mycoplasma* spp (Lookout Mycoplasma PCR Detection Kit, Sigma; St. Louis, MO). For some experiments, *C. trachomatis* serovar D (UW3-CX), *C. muridarum*, *C. caviae*, *C. pneumoniae* AR39, and *Coxiella burnetii* avirulent Nine Mile Phase II (provided by Bob Heinzen, Rocky Mountain Laboratories, Hamilton, MT) were used (11).

### Creation of Inc fusion constructs for transformation into *C. trachomatis* L2

All primers used in this study are listed in Table S4. All plasmids and *E. coli* strains used in cloning projects are listed in Table S5. The Inc-APEX2 fusion constructs were made as previously described (30). pcDNA3 APEX2-NES was a gift from Alice Ting (Addgene plasmid # 49386) (45). APEX2 contains a single N-terminal FLAG tag. For the construction of IncF_FLAG_, IncF with the C-terminal FLAG epitope was amplified from pTLR2 IncF-APEX2 (30) and cloned into pTLR2. CT226 was amplified from *C. trachomatis* L2 genomic DNA with primers containing a C-terminal FLAG tag and inserted into the mCherry site pBOMB4-Tet (EagI/KpnI) (a gift from Dr. T. Hackstadt, NIAID, Rocky Mountain Laboratories, Hamilton, MT) using NEBuilder HiFi Assembly Cloning Kit (NEBuilder). The final constructs were transformed into *dam^-^/dcm^-^ E. coli*. All constructs were confirmed by sequencing (Eurofins MWG Operon; Huntsville, AL).

### Transformation of *C. trachomatis* L2

Transformations were performed as described previously (89, 90). The Inc-APEX2 transformants (30) were plaque purified as described elsewhere (89, 91) and density gradient purified. Both pTLR2-IncF_FLAG_ and pBOMB4-Tet-CT226_FLAG_ were transformed as above in the presence of 1 U/mL penicillin and 1 μg/mL cycloheximide.

### Electron microscopy determination of APEX2 activity and localization

HeLa cells were seeded at 1.0 × 10^6^ cells/well in a 6-well plate containing 25 mm Thermanox cell culture treated coverslips for electron microscopy (Nunc; Rochester, NY). To confirm construct expression using indirect immunofluorescence, glass coverslips were included in duplicate wells of a 24-well plate. Wells were infected with *C. trachomatis* L2 wild-type, or L2 transformants, IncF-APEX2, IncA_TM_-APEX2, IncA-APEX2, or APEX2 only. *C. trachomatis* L2 IncF-APEX2, IncA_TM_-APEX2 (m.o.i. 0.75) were infected by centrifugation in DMEM + 10% FBS containing 2 U/mL penicillin and 1 μg/mL cycloheximide. *C. trachomatis* L2 IncA-APEX2 and APEX2 only (m.o.i. 0.75) infected wells received 1 U/mL penicillin and 1 μg/mL cycloheximide. *C. trachomatis* L2 wild-type was infected (m.o.i. of 0.4) in DMEM + 10% FBS containing 1 μg/mL cycloheximide. At 7 hpi, transformants were induced with 0.3 nM aTc for L2 IncF-APEX2, and 5 nM for all other L2 transformants and *C. trachomatis* L2 wild-type respectively. At 24 hpi, glass coverslips were fixed in 4% paraformaldehyde for 15 min at room temperature (RT) and methanol permeabilized for 5 min, then processed for immunofluorescence confirmation of construct expression as above.

The wells intended for electron microscopy were prepared using a protocol adapted from Martell et al. 2017 (44). Briefly, cells were washed with dPBS and fixing solution (2% glutaraldehyde, 2% paraformaldehyde in 0.1 M sodium cacodylate) was added to the wells and incubated on ice for 1 hour. All subsequent steps were performed on ice. The expressed APEX2 remains functional after fixing (using conditions below 4% formaldehyde). After 1 hour, the wells were washed 5 × 2 min with cold buffer A solution (0.1 M sodium cacodylate). To quench unreacted aldehyde groups, cold 20 mM glycine containing 2 mM CaCl_2_ in wash buffer A was incubated with the cells for 5 min. The wells were washed 5 × 2 min with cold buffer A. To enhance diffusion of the large molecule, 3,3′-Diaminobenzidine (DAB), the cells were pre-treated with 0.5 mg/mL DAB in buffer A containing 2 mM CaCl_2_ for 30 min prior to the polymerization step. The pre-treatment step allows the DAB to uniformly diffuse into the cells without being converted to the polymer (no H_2_O_2_ present). To catalyze the polymerization of DAB (regions proximal to APEX2) 0.5 mg/mL DAB and 3 mM H_2_O_2_ in buffer A containing 2 mM CaCl_2_ was added to cells and incubated for 30 min. Negative controls to determine background activity included *C. trachomatis* L2 wild-type with DAB treatment and *C. trachomatis* L2 IncA-APEX2, induced, without DAB. Polymerized DAB is unable to diffuse from the subcellular compartment. Finally, the wells were washed 5 × 2 min with cold buffer A and delivered to the University of Nebraska Medical Center electron microscopy core to be processed. In brief, the samples were post-fixed with 1% osmium tetroxide, stained with Toluidine Blue, dehydrated with a series of increasing ethanol concentrations, embedded and sectioned. Sections were placed on 200 mesh uncoated copper grids (Ted Pella Inc.), stained with uranyl acetate and Reynold’s lead citrate, and examined using a Tecnai G2 Spirit (FEI) transmission electron microscope (TEM) operated at 80 Kv. Representative electron micrographs are shown.

### FLAG affinity purification of APEX2 fusion constructs

HeLa cells were seeded in a 6-well plate in DMEM + 10% FBS and allowed to grow overnight. The cells were infected with *C. trachomatis* L2 IncF-APEX2, IncA_TM_-APEX2, IncA-APEX2, or APEX2 only (m.o.i. 0.75) with DMEM + 10% FBS containing 1 µg/mL cycloheximide, plus appropriate antibiotics (2 U/mL penicillin for *C. trachomatis* L2 IncF-APEX2 and IncA_TM_-APEX2, 1 U/mL penicillin for *C. trachomatis* L2 IncA-APEX2 and APEX2) and induced at 7 hpi with 0.3 nM aTc (IncF-APEX2 only; see (30) regarding lower induction levels), and 5 nM aTc (all other *C. trachomatis* L2 transformants). The cell collection, lysis procedure, and FLAG affinity purification were performed essentially as previously described (46). Briefly, at 24 hpi, coverslips were removed from the respective wells, methanol fixed for 5 min at RT, and the remaining cells were scraped into dPBS and centrifuged at 900 × g for 10 min at 4°C. The pellets were resuspended in cell lysis buffer [50 mM Tris-HCl, pH 7.4, 150 mM NaCl, 0.5% sodium deoxycholate, 0.1% sodium dodecyl sulfate (SDS), 1% Triton X-100 (Sigma; St. Louis, MO), 1X HALT protease inhibitor cocktail (Thermo; Waltham, MA), universal nuclease (Pierce; Rockford, IL), and 150 μM Clastolactacystin-β-lactone (Santa Cruz Biotechnology, Dallas, TX)]. Equal volumes (EZQ protein quantification; Life Technologies, Carlsbad, CA) of clarified lysates were prepared for FLAG affinity purification with FLAG magnetic beads (Sigma; St. Louis, MO) and rotated for 2 hours at 4° C. The affinity purified proteins were eluted in 30 µL of lysis buffer (above) containing FLAG peptide (200 µg/mL). The eluates from each sample were combined with an equal volume of 4x Laemmli sample buffer containing 5% β-mercaptoethanol and then loaded into a Criterion Midi 4-20% gradient SDS-PAGE (BioRad; Hercules, CA). The gel was transferred to PVDF (0.45 μm, Thermo Scientific; Waltham, MA) and blotted using anti-FLAG antibody to detect construct expression. Clarified lysate (used as the input for the FLAG affinity purification) was electrophoresed and transferred to PVDF to blot for chlamydial Hsp60 as a loading control.

### Labeling with biotin-phenol and affinity purification of biotinylated proteins

To identify proteins that were biotinylated using *C. trachomatis* L2 IncF-APEX2, IncA_TM_-APEX2, IncA-APEX2, and APEX2, HeLa cells were seeded into a 6-well plate in DMEM + 1% FBS. To monitor construct expression and biotinylation, coverslips were placed in 2 of the wells of the 6-well plate. The biotinylation assays were performed essentially as previously described (30-32). The cells were infected with *C. trachomatis* L2 IncF-APEX2, IncA_TM_-APEX2, IncA-APEX2, or APEX2 only (m.o.i. 0.75) with DMEM + 10% FBS containing 1 µg/mL cycloheximide, plus appropriate antibiotics (2 U/mL penicillin for *C. trachomatis* L2 IncF-APEX2 and IncA_TM_-APEX2, 1 U/mL penicillin for *C. trachomatis* L2 IncA-APEX2 and APEX2), and centrifuged 400 × g at RT for 15 min. Penicillin and cycloheximide were present for all biotinylation experiments to preserve the integrity of our transformants and to minimize host cell background, respectively. The samples were induced for construct expression at 7 hpi with 0.3 nM aTc (IncF-APEX2 only; see (30) regarding lower induction levels) or 4 nM aTc (all other transformants). At 23.5 hpi, the cell monolayers were incubated with 1.5 mM biotinyl-tyramide (biotin-phenol) (AdipoGen, San Diego, CA) for 30 min at 37°C + 5% CO_2_. At 24 hpi, the labeling process was catalyzed, quenched, and the lysate was collected as previously described (30). Normalized lysates were added to equilibrated streptavidin magnetic beads (Pierce; Rockford, IL) and rotated for 90 min at RT. Proteins were eluted from streptavidin magnetic beads by 4 min incubation at 95 °C in 2x Laemmli sample buffer containing 0.5 mM biotin. The eluates were loaded into Criterion Midi 4-20% gradient denaturing gels (BioRad; Hercules, CA) in duplicate. The gel intended for Coomassie staining was resolved briefly (~2-3 cm), then stained (10% methanol, 5% acetic acid, Coomassie blue G). The duplicate gel, for western blot confirmation of affinity purification was resolved completely, transferred to PVDF (0.45 μm, Thermo Scientific; Waltham, MA) and blotted using the indicated primary antibodies, a streptavidin-488 conjugate (immunofluorescence) or streptavidin-680 conjugate (western blot), and appropriate secondary antibodies conjugated to IRDye 680LT, IRDye CW, or a streptavidin-IRDye 680LT conjugate (LiCor Biosciences, Lincoln, NE). The PVDF membranes were imaged using an Azure c600 (Azure Biosystems, Dublin, CA) and processed using Adobe photoshop creative cloud (Adobe).

#### Identification of biotinylated proteins using mass spectrometry

Coomassie-stained gels were imaged, then each lane was cut into six gel fractions to enhance the resolution of lower abundance proteins. The UNMC proteomics core facility performed in-gel digestion, preparation, and analysis of gel fractions. Protein fractions excised from the SDS-PAGE were destained, reduced with tris-carboxyethyl phosphine, alkylated with iodoacetamide, and were digested overnight with sequencing-grade trypsin (Promega; Madison, WI) and Asp-N (Promega; Madison, WI). Trypsin (cleaves Lys and Arg) and Asp-N endoproteinase (cleaves Asp and Cys residues). Peptides were eluted from the gel and concentrated to 20 μL by vacuum centrifugation and analyzed using a high-resolution mass spectrometry nano-LC-MS/MS Tribrid system, Orbitrap Fusion™ Lumos™ coupled with UltiMate 3000 HPLC system (Thermo Scientific; Waltham, MA). Approximately 500 ng of peptides were run by the pre-column (Acclaim PepMap™ 100, 75μm × 2cm, nanoViper, Thermo Scientific; Waltham, MA) and the analytical column (Acclaim PepMap™ RSCL, 75 μm × 50 cm, nanoViper, Thermo Scientific; Waltham, MA). The samples were eluted using a 100-min linear gradient of Acetonitrile (2.5-45 %) in 0.1% Formic acid.

All MS/MS samples were analyzed using Mascot (Matrix Sciences, London, UK, vs. 2.6.). Mascot was set up to search the SwissProt database (selected for Homo sapiens, 2017_02, 20286 entries and *C. trachomatis* 434/Bu entries) assuming the digestion enzymes trypsin and AspN. Parameters on MASCOT were set as follows: Enzyme: Trypsin and Asp-N for biological replicates n=5, Max missed cleavage: 2, Peptide charge: 1+, 2+ and 3+, Peptide tolerance: ± 0.8 Da, Fixed modifications: carbamidomethyl (C), Variable modifications: oxidation (M) and biotin-phenol (C, Y, W, H). MS/MS tolerance: ± 0.6 Da, Instrument: ESI-TRAP. Proteins identified by Mascot search were uploaded into Scaffold for visualization of the identified proteins (Scaffold, Proteome Software, Inc. Portland, Oregon).

### Statistical analysis of mass spectrometry samples using Significance Analysis of INTeractome (SAINT)

Significance Analysis of INTeractome (SAINT) was performed to assign statistical significance (Bayesian false discovery rate; BFDR) to our mass spectrometry data (38). SAINT calculates the probability that a protein identified in the test sample is a true interacting protein based on average hits in the test samples compared to the control in an unbiased fashion. Scaffold files containing each replicate (n=5) were set to 95% protein threshold, 1 peptide minimum, and 95% peptide threshold, and the sample report was exported to an excel file. The sample report file was used to make three files required for SAINT analysis: bait, prey, and interaction (Table S1 and S3). The bait file corresponds to the sample condition (e.g., IncF-APEX2, replicate 1, Test condition) and assigns samples as either a test or control. For our dataset, the Inc-APEX2 biotinylated proteins via IncF-APEX2, IncA_TM_-APEX2, IncA-APEX2 are the test, “T”, and the controls “C” were assigned to APEX2, L2 wild-type, and mock-infected HeLa cells. The prey file is the list of all the proteins from the Scaffold sample reports file using their gene names and amino acid length (obtained from UniProt). The last file required for SAINT is the interaction file which assigns the biological replicate number and spectral counts for each protein identified in the test subjects and the control samples. These files are input to calculate the Bayesian False Discovery Rate (BFDR) and were used to prioritize which proteins were statistically significant (38). We then input high confidence data (BFDR ≤0.05) into the pipeline to visualize interaction networks created using the STRING database (interaction confidence 0.7 STRING). The defined STRING networks were exported and analyzed using Cytoscape v 3.7.1 (51) with ClueGo to determine globally enriched biological processes and molecular functions within each dataset.

### Transfection of LRRF1-GFP and FLI1-GFP

LRRF1 detected by mass spectrometry corresponded to LRRF1 variant 3. To assess LRRF1 and FLI1 localization during *C. trachomatis* L2 infection, we obtained pCMV6-AC-LRRF1-GFP (LRRF1 variant 3; origene #RG226542; Rockville, MD) and pCMV6-AC-FLI1-GFP (Origene # RG206863; Rockville, MD) respectively. For DNA transfections, 8 × 10^4^ HeLa cells per well were seeded in a 24-well plate onto 12 mm glass coverslips. Approximately 24 hours later fresh DMEM + 10 % FBS (antibiotic free) was added to the cells. Transfection efficiency was first optimized using varying nanogram amounts of pDNA and volumes of jetPRIME® transfection reagent (Polyplus; Illkirch, France). Optimal efficiency was determined at 100 ng of pCMV-LRRF1-GFP or 500 ng pCMV6-AC-FLI1-GFP added to 50 µL of jetPRIME® buffer and 1.0 µL of transfection reagent (Polyplus; Illkirch, France). Samples were vortexed for 10 sec, centrifuged briefly, and incubated at RT for 10 min. The plasmid/transfection reagent mixture was added dropwise to individual wells. After four hours post-transfection, the media was changed and two hours later (6 hours post-transfection), HeLa cells were infected with *C. trachomatis* L2 wild-type (m.o.i. 0.8) by centrifugation at 400 × g for 15 min at RT. At 24 hpi, cells were fixed in 4% paraformaldehyde, permeabilized with 0.5 % Triton X-100 for 5 min at RT and stained for immunofluorescence to visualize the inclusion membrane (anti-CT223), LRRF1-GFP or FLI1-GFP, and DNA (DAPI). The coverslips were imaged using Zeiss with Apotome.2 at 100x. Scale bar = 10 µm.

### siRNA knockdown of LRRF1 to determine the effect on infectious progeny production

siRNA knockdown experiments were performed following the manufacturer’s protocol (Polyplus; Illkirch, France). Non-targeting siRNA (Origene #: SR30004; Rockville, MD), GAPDH siRNA (Ambion cat #4390849), and pooled LRRF1 siRNA (Ambion Life Technologies siRNA cat #43450, s229968, and s17599) were used in knockdown experiments. siRNA experiments were set up in quadruplicate to confirm LRRF1 knockdown efficiency by western blot (one well), to detect LRRF1 (unpublished) or FLI1 localization by immunofluorescence (one well), and to quantify infectious progeny (two wells). Briefly, 20 nM of the non-targeting (NT), GAPDH, single LRRF1 siRNA, or pooled LRRF1 siRNA was added to serum free Opti-MEM (100 μL/well), and 2 μL of INTERFERin reagent (Polyplus; Illkirch, France) was added to each well. The wells were incubated for 15 min at RT with gentle rocking. Then 2.5 × 10^4^ HeLa cells were added to each well on top of the siRNA/transfection reagent mixture and incubated at 37 °C + 5% CO_2_. The media was replaced with fresh DMEM + 10% FBS after 24 hours. At 48 hours post-siRNA transfection, HeLa cells were infected with *C. trachomatis* L2 wild-type (m.o.i. 0.8) by centrifugation at 400 × g 15 min at RT.

At 30 hpi, to confirm knockdown efficiency, *C. trachomatis* L2 infected HeLa cells were trypsinized, centrifuged and resuspended in 2x Laemmli sample buffer. The lysate was loaded, electrophoresed, and transferred to PVDF, then blotted to detect the presence of LRRF1 and GAPDH. To measure infectious progeny, experiments were performed essentially as previously described (92, 93). Briefly, duplicate wells for each sample were lysed at 30 hpi by scraping and then centrifuged at 17,000 × g for 30 min at 4 °C. The pellet was resuspended in sucrose phosphate buffer (2SP), serially diluted, and infected in duplicate onto a fresh monolayer of HeLa cells by centrifugation at 400 × g for 15 min at RT. Cells were incubated at 37°C + 5% CO_2_ for 15 min, then the 2SP buffer was replaced with DMEM + 10 % FBS containing 1 μg/mL cycloheximide. To quantify infectious progeny, at 24-30 hours post-secondary infection, the cells were fixed in methanol for 5 min at RT and processed for indirect immunofluorescence to visualize the inclusion using anti-MOMP antibodies (Meridian Biosciences; Memphis, TN). The average inclusion forming units (IFU/mL) from three biological replicates is reported.

### Validation of LRRF1 at the chlamydial inclusion and time course experiments

HeLa cells infected with *C. trachomatis* L2 (m.o.i 0.75) in DMEM + 10% FBS without antibiotics were fixed at 24 hpi in 4% paraformaldehyde, permeabilized with 0.5 % Triton X-100 for 5 min at RT and stained for immunofluorescence to visualize the inclusion membrane (anti-CT223), LRRF1, and DNA (DAPI). The coverslips were imaged using Zeiss with Apotome.2 at 100x. Scale bar = 10 µm.

For the time course experiments, HeLa cells infected with *C. trachomatis* L2 (m.o.i 0.75) or mock-infected in DMEM + 10% FBS without antibiotics were fixed at 8, 12, 16, 24, and 36 hpi in methanol for 5 min at RT. One sample was treated with 34 µg/mL chloramphenicol at 8 hpi and then methanol fixed at 36 hpi. Fixed coverslips were stained for immunofluorescence to visualize the inclusion membrane (anti-CT223), LRRF1, *Chlamydiae* (MOMP), and DNA (DAPI) and imaged using Zeiss with Apotome.2 at 100x. Scale bar = 10 µm.

### Assessing LRRF1 localization during infection of *C. trachomatis* serovars, *Chlamydia* species and *Coxiella burnetii*

HeLa cells infected with *C. trachomatis* L2 (m.o.i 0.75), *C. trachomatis* serovar D (m.o.i 1), *C. muridarum* (m.o.i 0.25), *C. caviae* (m.o.i 0.25), *C. pneumoniae* (m.o.i 1), and *Coxiella burnetii* avirulent Nine Mile Phase II were used. DMEM + 10% FBS media did not contain antibiotics (penicillin or cycloheximide) for these experiments. *C. trachomatis* serovar D was pre-treated with DEAE-Dextran prior to infection. All *Chlamydia*-infected HeLa cells were centrifuged at 400 × g 15 min at RT. *C. burnetii* Nine Mile Phase II infected HeLa cells were centrifuged for 1 hr at 2000 rpm at RT. At 24 hpi, *C. trachomatis* L2, *C. trachomatis* serovar D, *C. muridarum*, and *C. caviae* infected HeLa cells were methanol fixed and stained for immunofluorescence. At 96 hpi, *C. pneumoniae* infected HeLa cells were fixed in 4% paraformaldehyde, permeabilized with 0.5% Triton X-100 and stained for immunofluorescence. At 3 days post infection, *C. burnetii* Nine Mile Phase II infected HeLa cells were methanol fixed for 5 min at RT and stained. Coverslips were stained using organism-specific and LRRF1 antibodies listed in to examine LRRF1 localization and DRAQ5 or DAPI to visualize DNA.

### Bacterial adenylate cyclase two-hybrid (BACTH) assays

To screen for LRRF1 interacting partners, *LRRF1* was amplified from the pCMV-LRRF1-GFP vector (Origene; Rockville, MD), and *Incs* were amplified from *C. trachomatis* L2 genomic DNA using primers with overlapping sequences for each pST25 and pUT18C vectors (Table S4, Table S5). *LRRF1*, *CT288, CT226, CT223, IncA, IncF* amplified using the primers listed in Table S5 were cloned into each either pST25 or pUT18C using the NEbuilder HiFi Assembly Cloning Kit (NEBuilder) and transformed into DH5α lacI^q^ *E. coli*. Individual clones were cultured overnight, pDNA was isolated (Qiagen; Germantown, MD), verified by restriction digest, and the final clones were verified by DNA sequencing. pUT18C-IncF (Gateway®) (serovar D) was made as previously described (23). To screen for interactions, assays were performed as described previously (23, 64, 67, 71, 94). Briefly, plasmids were co-transformed into DHT1 (Δ*cyaA*) *E. coli* (Table S4) and prior to plating, *E. coli* cells were pelleted, washed and resuspended in 1x M63 minimal media. The resuspended DHT1 *E. coli* were then plated on 1x M63 minimal media plates containing, 0.2% Maltose, Isopropyl β-D-1-thiogalactopyranoside (IPTG; 0.5 mM), 5-bromo-4-chloro-3-indolyl-β-D-galactopyranoside (X-gal; 0.04 mg/mL), casamino acids (0.04%), spectinomycin (25 µg/mL), and ampicillin (50 µg/mL) and incubated at 30 °C for 3-5 days. To quantify interactions by β-galactosidase assay, eight colonies (or streaks from negative plates) were set up for overnight culture in minimal media (1x M63 containing 0.2% maltose, IPTG 0.5 mM, 0.04 mg/mL X-gal, 0.01% casamino acids, spectinomycin (25 µg/mL), and ampicillin (50 µg/mL). After 20-24 hours, the cultures were diluted, and the OD600 was measured. A duplicate set of samples were permeabilized with SDS (0.05 %) and chloroform. After permeabilization, the supernatant was transferred to an optical plate containing 0.4% ONPG in PM2 buffer with 100 mM 2-mercaptoethanol. After 20 minutes, the enzymatic reaction was stopped with 1 M Na_2_CO_3_ stop solution and the absorbance at 405 nm was measured. The OD405 was normalized to bacterial growth (OD600) and reported as relative units (RU). A positive interaction is defined as greater than five times the negative control (66). Three independent experiments were analyzed for each interaction and graphed using GraphPad Prism 7 and reported as the mean with standard deviation.

### Super resolution microscopy to assess LRRF localization with Incs

HeLa cells seeded on glass coverslips were infected with *C. trachomatis* L2 wild-type, or *C. trachomatis* L2 transformants, IncF-APEX2, IncA_TM_-APEX2, IncA-APEX2, CT226_FLAG_ and induced for expression at 20 hpi (5 nM aTc for all transformants except IncF-APEX2 was induced with 1 nM aTc). At 24 hpi, coverslips were rinsed once with dPBS and then fixed with ice cold methanol and stained for immunofluorescence to visualize construct expression (FLAG) or CT223 (red), LRRF1 (green), *Chlamydiae* and DNA (blue). Coverslips were imaged using Zeiss ELYRA PS.1 Super Resolution Microscope Zeiss with Structured Illumination Microscopy (SIM). Scale bar = 5 µm. Using Zen Blue (Zeiss), 3D snapshots from *C. trachomatis* L2 CT226_FLAG_ infected HeLa cells and *C. trachomatis* L2 IncA-APEX2 infected HeLa cells with IncA fibers were generated and exported for visualization.

### Co-immunoprecipitation of CT226_FLAG_ with endogenous LRRF1

HeLa cells were seeded in a 6-well plate in DMEM + 10% FBS and allowed to grow overnight. A coverslip was placed in two wells of each 6-well plate to monitor construct expression and localization by indirect immunofluorescence for each experiment. The cells were infected with *C. trachomatis* L2 CT226_FLAG_ and IncF_FLAG_ (m.o.i. 0.8) with DMEM + 10% FBS containing 1 U/mL penicillin (no cycloheximide) and not induced or induced with 5 nM (CT226_FLAG_) or 1 nM aTc (IncF_FLAG_) at 7 hpi. At 24 hpi, coverslips were removed, fixed in 4% paraformaldehyde, triton-X permeabilized (0.5%) and stained for immunofluorescence to detect construct expression (FLAG), the inclusion membrane (IncA), DNA and *Chlamydiae*. The cells were lysed and affinity purified using FLAG magnetic beads as described above and previously (46). The eluates were mixed with an equal volume of 4x Laemmli sample buffer containing 5% β-mercaptoethanol and then loaded into a Criterion Midi 4-20% gradient SDS-PAGE (BioRad; Hercules, CA). The gel was transferred to PVDF (0.45 μm, Thermo Scientific; Waltham, MA) and blotted using anti-FLAG antibody to detect construct expression and anti-LRRF1 antibody. Three biological replicates were analyzed by co-immunoprecipitation.

## Acknowledgments

We would like to thank R. Suchland (University of Washington, WA) and Dr. D. Rockey (Oregon State University, OR) for the anti-CT223 antibody, Dr. T. Hackstadt (NIAID, Rocky Mountain Laboratories, Hamilton, MT) for the anti-IncA antibody, Dr. R. Morrison (University of Arkansas for Medical Sciences, Little Rock, AR) for the anti-chlamydial Hsp60 antibody, and Dr. H. Caldwell (NIAID, Bethesda, MD) for the *C. pneumoniae* anti-MOMP antibody. We would also like to thank Dr. Bob Heinzen (NIAID, Rocky Mountain Laboratories, Hamilton, MT) for avirulent *Coxiella burnetii* Nine Mile Phase II strains.

The authors would also like to thank Eric Troudt and Vikas Kumar for processing samples for mass spectrometry and their technical expertise, and Tom Bargar for processing samples for electron microscopy. The transmission electron microscope is supported by NIH Shared Instrument grant (NIH 1 S10 RR024650 01A1). This work was also supported by the UNMC Center Advanced Microscopy Core Facility and the UNMC Advanced Proteomics Core Facility, and the UNMC Electron Microscopy Core Facility. The University of Nebraska Medical Center Advanced Microscopy Core Facility receives partial support from the National Institute for General Medical Science (NIGMS) INBRE - P20 GM103427 and COBRE - P30 GM106397 grants, as well as support from the National Cancer Institute (NCI) for The Fred & Pamela Buffett Cancer Center Support Grant - P30 CA036727, and the Nebraska Research Initiative. This publication’s contents and interpretations are the sole responsibility of the authors.

## Supporting information

**S1 Fig. Western blot detection of the expressed APEX2 constructs.**

HeLa cells infected with *C. trachomatis* L2 transformants or wild-type (WT) were induced with anhydrotetracycline (aTc) at 7 hpi (0.3 nM for IncF-APEX2; all other samples 5 nM), or uninduced as indicated. Lysates were collected at 24 hpi, solubilized, and affinity purified using FLAG beads. The eluates were separated by electrophoresis, transferred to PVDF membrane, and blotted for APEX2 containing constructs using anti-FLAG antibody. The total lysate (input) was probed with anti-*C. trachomatis* Hsp60 (cHsp60) antibody as a loading control.

**S2 Fig. Visualization of global biological processes and molecular function of AP-MS identified statistically significant eukaryotic proteins using *C. trachomatis* L2 Inc-APEX2 transformants.**

Affinity purified-mass spectrometry (AP-MS) identified spectra were compared to the *Homo sapiens* database using Mascot and then Significance Analysis of INTeractome (SAINT) was applied to identify statistically significant proteins (BFDR ≤ 0.05) from each dataset. Global networks identified using each *C. trachomatis* L2 transformed with (A) IncF-APEX2, (B) IncA_TM_-APEX2, and (C) IncA-APEX2 are shown.

**S3 Fig. STRING network analysis of statistically significant eukaryotic proteins.**

Significance Analysis of INTeractome (SAINT) was applied to identify statistically significant AP-MS identified eukaryotic proteins using *C. trachomatis* L2 IncF-APEX2, IncA_TM_-APEX2, and IncA-APEX2. STRING network (0.7 high confidence) visualization of eukaryotic proteins identified by mass spectrometry (SAINT BFDR ≤ 0.05) from each *C. trachomatis* L2 (A) IncF-APEX2, (B) IncA_TM_-APEX2, and (C) IncA-APEX2.

**S4 Fig. The effect of LRRF1-GFP and FLI1-GFP overexpression and LRRF1 knockdown on *C. trachomatis* progeny production.**

HeLa cells seeded onto coverslips were transfected with (A) 100 ng pCMV6-AC-LRRF1-GFP or (B) 500 ng pCMV6-AC-FLI1-GFP. Transfected cells were either mock-infected or infected with *C. trachomatis* L2 wild-type at 6 hours post-transfection. At 24 hpi, HeLa cells were paraformaldehyde fixed 0.5 % Triton X-100 permeabilized and stained for immunofluorescence to visualize the inclusion membrane (CT223; red), DNA (DRAQ5; blue) and (A) LRRF1-GFP or (B) FLI1-GFP. Coverslips were imaged using a Zeiss with ApoTome.2 at 100x. Scale bar = 10 µm. (C) 20 nM of non-targeting (NT), GAPDH, LRRF1 single siRNA or 3 pooled siRNAs were reverse transfected as indicated into HeLa cells that were then infected with *C. trachomatis* L2 wild-type at 48 hours post-transfection and collected 24 hours later. Lysates from siRNA treated, *C. trachomatis* L2 wild-type infected cells were collected in 2x Laemmli sample buffer, electrophoresed, transferred to PVDF and blotted to confirm siRNA knockdown efficiency of LRRF1 and GAPDH. Duplicate wells were lysed, infected onto a fresh HeLa cell monolayer and enumerated for progeny production. Three independent experiments were performed and averaged (Non-targeting siRNA= 2.73 × 10^6^ IFU/mL; GAPDH siRNA=4.46 × 10^6^ IFU/mL; Single LRRF1 siRNA= 2.3 × 10^6^ IFU/mL; Pooled LRRF1 siRNA= 4.09 × 10^6^ IFU/mL).

**S5 Fig. Co-immunoprecipitation of endogenous LRRF1 with *C. trachomatis* L2 CT226_FLAG_**

HeLa cells seeded in a 6-well plate with glass coverslips were infected with *C. trachomatis* L2 CT226_FLAG_ or IncF_FLAG_ and either not induced or induced for expression at 7 hpi with 5 nM aTc (CT226_FLAG_) or 1 nM aTc (IncF_FLAG_). At 24 hpi, cells were collected, solubilized, normalized, and affinity purified using FLAG beads. The clarified lysates (soluble), unbound fractions and eluates were probed for construct expression (FLAG; CT226_FLAG_ 19.2 kDa; IncF_FLAG_ 11.3 kDa monomer and 22.6 kDa dimer) and LRRF1 (dimer 160 kDa). (A) and (B) are representative of two biological replicates. A total of three independent experiments were performed.

**Table S1. Complete SAINT analysis of *C. trachomatis* L2 proteins identified by mass spectrometry**

The completed Mascot database search files were uploaded into Scaffold (v 4.8.7) and the samples report containing protein identification and spectral abundance data was exported to excel. Three files (bait, prey, interaction) were created for input into the SAINT program. The SAINT results file organized by Bayesian False Discovery Rate (BFDR). The bait proteins are color coded, and the statistically significant proteins (BFDR ≤0.05) are highlighted.

**Table S2. SAINT significant eukaryotic proteins identified by mass spectrometry**

SAINT analyzed statistically significant eukaryotic proteins organized by bait (Inc-APEX2) sample and Bayesian False Discovery Rate (BFDR ≤0.05). The highlighted proteins were identified as statistically significant using all three bait (Inc-APEX2) constructs.

**Table S3. Complete SAINT analysis of eukaryotic proteins identified by mass spectrometry**

The completed Mascot database search files were uploaded into Scaffold (v 4.8.7) and the samples report containing protein identification and spectral abundance data was exported to excel. Three files (bait, prey, interaction) were created for input into the SAINT program. The SAINT results file organized by Bayesian False Discovery Rate (BFDR). Boosted by Corum indicates proteins that are known to interact with the prey protein and were also detected in this dataset. The bait proteins are color coded, and the statistically significant proteins (BFDR ≤0.05) are highlighted.

**Table S4. *E. coli* and plasmids used in this study**

**Table S5. Primers used in this study**

